# Mosaic display of stable hemagglutinin monomers induces broad immune responses

**DOI:** 10.1101/2024.04.10.588945

**Authors:** Hyojin Kim, Seong Cheol Min, Dan Bi Lee, Ju Hwan Jeong, Eun Jeong Kim, Jin Young Maeng, Jong Hyeon Seok, Ji-Hye Lee, Yuyeon Jang, Ui Soon Jang, Tine Ysenbaert, João Paulo Portela Catani, JeongAh Lee, Yeongeun Lee, Youn Kyung Kim, Gyudo Lee, Ji Young Mun, Hyun Sik Jun, Yun Hee Baek, Xavier Saelens, Jiwon Lee, Mi Sook Chung, Min-Suk Song, Kyung Hyun Kim

## Abstract

The nature of the interplay between immunity and viral variation is infinitely adaptive. Infection frequently induces immune responses against variation-prone epitopes, rather than against spatially hidden conserved epitopes. It thus remains a substantial challenge to elicit the immune responses to the conserved epitopes providing broad-spectrum immunity. We developed an approach of scaffold-mediated mosaic display to present monomeric influenza virus hemagglutinins (HAs), which exposes highly conserved stem and interface epitopes. Stable monomers were rationally engineered from H1 and H3 subtypes and B type HA trimers, with amino acid mutations at the monomer-monomer interface and for disulfide bond formation, and fused to a self-assembling scaffold, to generate a mosaic HA monomer-displaying nanoparticle, 3HA-np. Immunization with 3HA-np induced broadly neutralizing antibodies (bnAbs) in mice and ferrets and protected against challenges with H1N1 and H3N2 viruses. Competitive immunoassays revealed that 3HA-np induced high interface- and stem-binding Ab titers as compared to head Ab titers, indicating that the monomeric and mosaic nature of 3HA-np elicit cross-reactive Abs. Our results suggest that exposure of the hidden conserved epitope by monomer-displaying nanoparticles is a promising approach to generate a universal influenza vaccine.

## INTRODUCTION

Influenza A and B viruses cause seasonal flu epidemics in humans. Influenza A subtypes (H1N1 and H3N2) and influenza B lineages (Yamagata and Victoria) that are antigenically distinct are currently circulating, although the Yamagata lineage has not been isolated since the start of the COVID-19 pandemic^1^. Human influenza A and B viruses change rapidly through antigenic drift, which causes frequent antigenic mismatch between the circulating and vaccine strains. Antigenically distinct strains arise sporadically through antigenic shift, leading to global influenza pandemics as seen in 1957 (H2N2), 1968 (H3N2) and 2009 (H1N1), which are partly of avian-origin^2,3^. Seasonal influenza vaccines have shown variable effectiveness against diseases caused by circulating strains, despite yearly updates^4^, and even less protection against antigenically shifted influenza A viruses, such as H5N1 and H7N9, which can cause zoonotic infection^5^.

The active form of hemagglutinin (HA) is composed of two distinct regions: the head region with the receptor binding domain and the stem region with long central helices. Unlike the variable and immunodominant head, the immunosubdominant stem shows a high degree of sequence conservation. It undergoes pH-induced conformational changes that result in membrane fusion, a key step in the influenza virus entry process^6,7^. While seasonal vaccines primarily induce strain-specific neutralizing antibodies (nAb) targeting the variable head region of HA, necessitating near annual updates, universal influenza vaccines should be protective against antigenically diverse influenza A and B viruses^8^. Immunization of mice with HA-ferritin nanoparticles demonstrated enhanced breadth of coverage against unmatched viruses^9^. Directing the induction of immune responses toward the conserved stem, referred to as immunofocusing, led to the identification of non-neutralizing but cross-reactive Abs (crAbs)^10,11^. The stem attached to virus-like particles further revealed broad cross-reactivity across group 1 viruses^12−15^. Phase I clinical trials of a chimeric HA or stem on nanoparticles induced broadly neutralizing immune responses^16,17^.

Molecular characterization of serum Ab repertoires revealed a prevalence of crAbs that bind to the conserved monomer-monomer interface, occluded inside the HA trimer^18^. The interface-specific Abs conferred full cross-protection *in vivo* and were able to dissociate HA trimers into monomers upon binding *in vitro*^19−21^. The observation is seemingly contrary to that of conventional epitope binding and neutralization that would require HA trimerization to attain full antigenicity. However, monomeric HA-binding Abs show binding affinity comparable to that of trimer-binding Abs^22^. We showed previously that the recombinant HA from a pandemic strain A/Korea/01/2009 was monomeric in solution^23^, and that a stable monomer derived from the A/Thailand/CU44/2006 HA trimer, by mutations at the monomer-monomer interface and for disulfide bond formation, induced *in vivo* protective immunity, comparable to the trimer^24^. Despite the identification of interface or stem-binding crAbs, how to elicit humoral immune responses to the conserved epitopes remains a substantial challenge associated with inducing broad-spectrum immunity.

Co-display of heterotypic HA on nanoparticles, by designed scaffold or Spy-tag-mediated fusion chemistry, elicited broader Ab responses than a mixture of homotypic nanoparticles or commercial tri- or quadrivalent vaccines^25,26^. The mosaic nanoparticles confer an avidity advantage to cross-reactive B cells, whereas homotypic antigens or multiple boosters typically promotes the induction of strain-specific B cells. However, current mosaic nanoparticle platforms suffer from a major drawback of non-uniform antigen assembly, primarily due to problems to produce nanoparticles that uniformly incorporate heterotypic antigens. Here, we explored an approach of mosaic nanoparticles that incorporate stable H1, H3, and B HA monomers into a self-assembling scaffold, the proliferating cell nuclear antigen (PCNA) from the archaeal species *Saccharolobus solfataricus*^27^. The archaeal PCNA scaffold comprise three subunits of PCNA1, PCNA2 and PCNA3, which form a ring-shaped heterotrimer, whereas PCNA from bacteria to higher animals including humans are homotrimeric^28−30^. The mosaic nanoparticle, 3HA-np described here, presents the diverse HA monomers uniformly and elicited heterosubtypic and heterotypic Ab responses and could protect mice and ferrets against both H1N1 and H3N2 virus challenges.

## RESULTS

### Design and assembly of mosaic nanoparticle 3HA-np

Starting from the HA trimers derived from A/California/04/2009 (H1N1) (CA04), B/Florida/4/2006 (FL04), A/Gyeongnam/684/2006 (H3N2) (Gy684), five to eight amino acid residues at the monomer-monomer interface were mutated to stabilize the HA monomers and prevent their association into trimers (Fig. 1A & Supplementary Fig. S1 & S2). In the cases of H1 and H3 HAs, mutations were introduced to allow disulfide bond formations to further stabilize the HA monomers. The HA monomers engineered by the interface mutations and disulfide bond formations were expressed, purified by His-tag affinity and size exclusion chromatography (SEC), and assessed by SEC-multiangle light scattering (SEC-MALS) and differential scanning fluorimetry (DSF) (Fig. 1B). H1 HA_m_, B HA_m_, and H3 HA_m_ had molecular weights of 62.5–85.0 kDa. The mutations were designed to reduce intermonomer interactions and to cause hindrance upon trimerization of HA, by electrostatic or steric repulsion, where negatively charged or large amino acids are positioned very close at the stem region. Nevertheless, the mutant monomers showed slightly less stability than the corresponding trimers, with lower melting temperature T_m_ (1.0–3.6°C), except for the H1 HA_m_, which showed higher Tm (∼8.5°C) than the recombinant H1(CA04) HA that was monomeric in solution^23^ (Fig. 1B).

**FIGURE 1.**
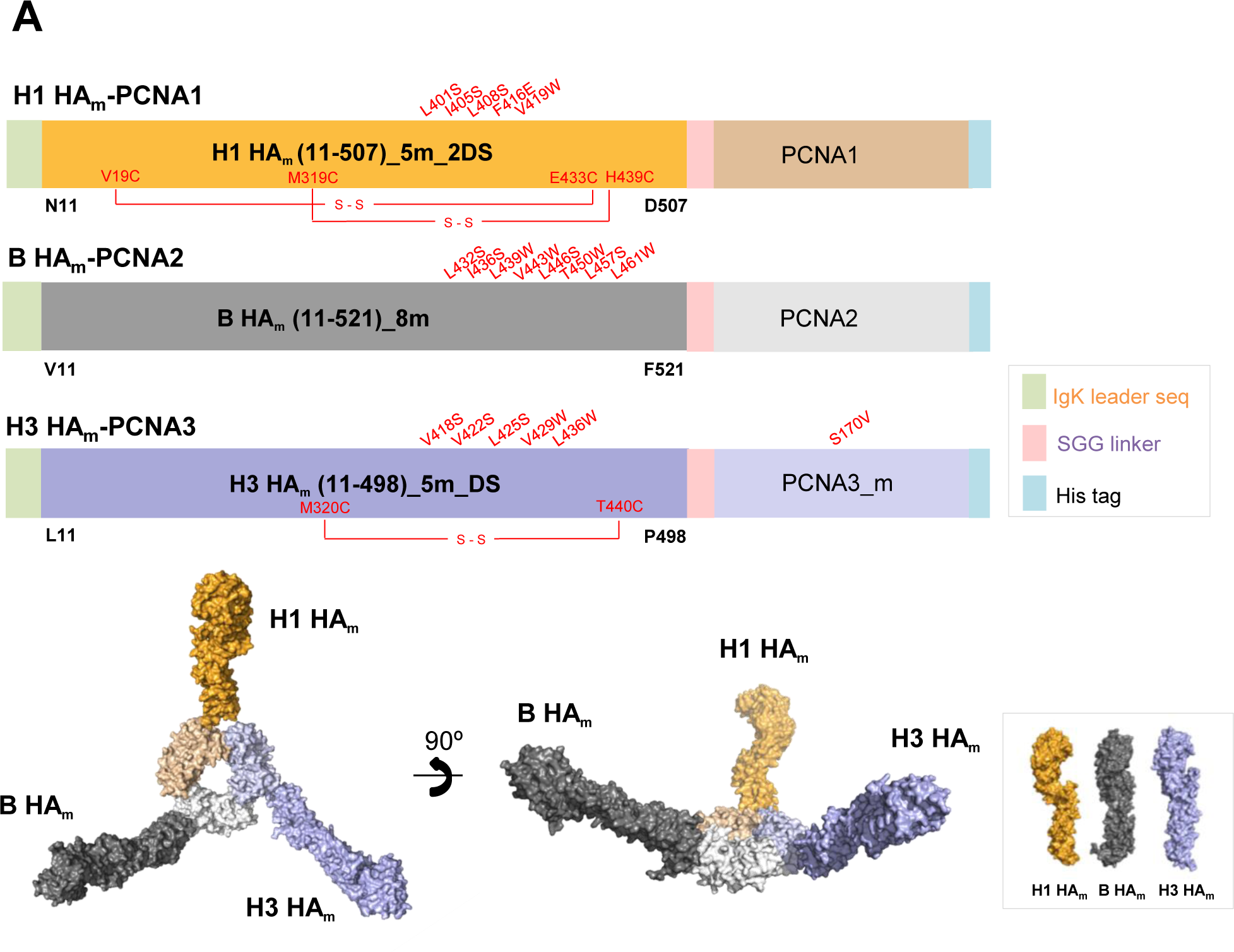
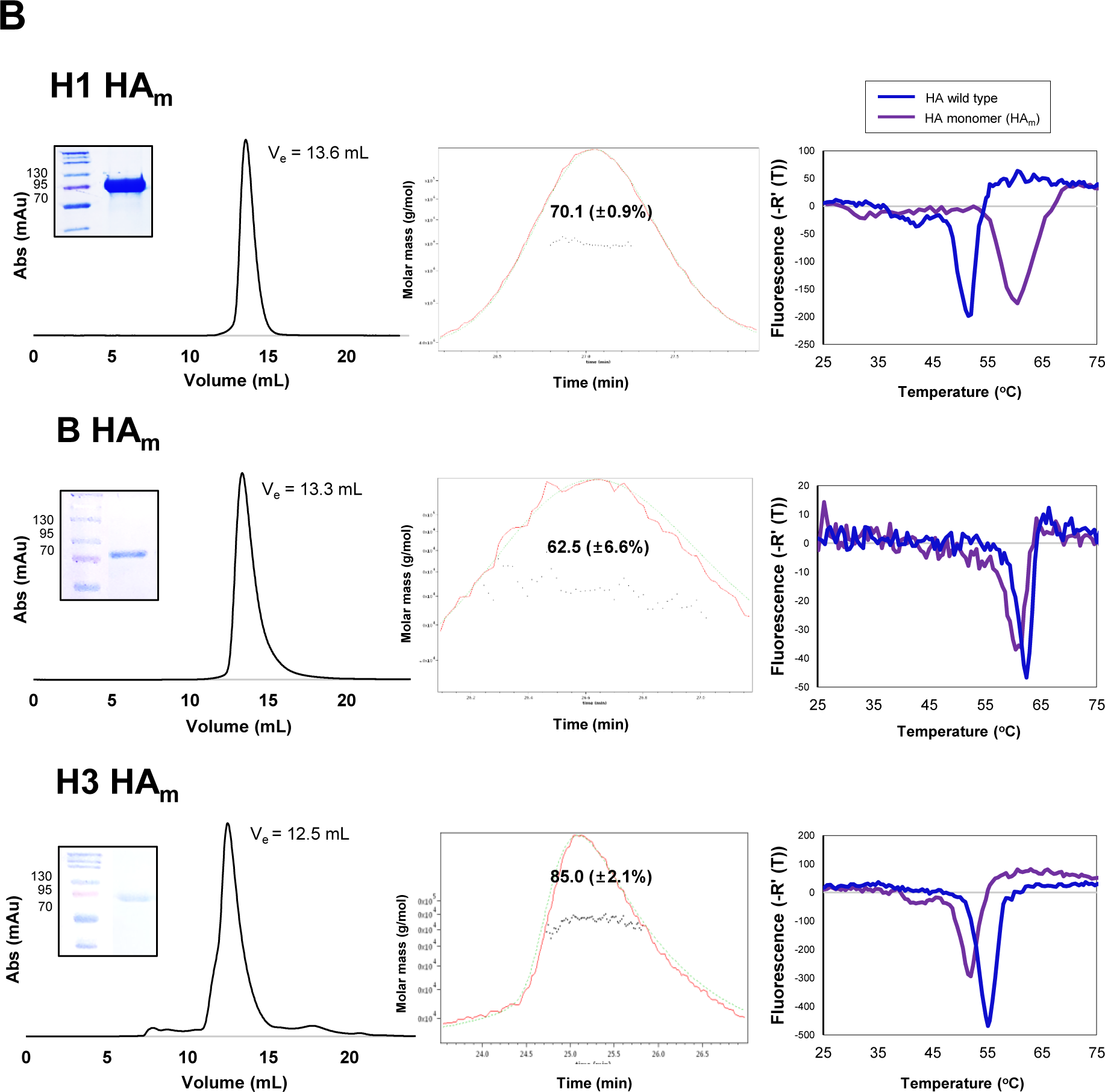
Design and characterization of antigens. (A) Design of H1 HA_m_-PCNA1, B HA_m_-PCNA2, and H3 HA_m_-PCNA3 showing schematic diagrams of influenza virus HA-PCNA fusion proteins. The HA regions are derived from 11 to 507 for H1, 11 to 521 for B, and 11 to 498 for H3 HA proteins, where mutation sites for stable HA monomer (HA_m_) are indicated by red texts. The HA is fused to the N-terminus of PCNA via SGG linker, and the His-tag is fused to C-terminus of the gene for mammalian cell expression (upper panel). A molecular model of 3HA-np displaying PCNA ring and 3HA_m_ proteins colored as in the schematic diagram (lower panel), showing that the HA_m_s fused to the PCNA subunits are exposed on the external surface of the nanoparticle. The models were generated from structures of PDB IDs: 4EDB, 4M40, 2YP7, and 2HIK for H1, B, H3 HA proteins, and PCNA, respectively. (B) Characterization of H1 HA_m_, B HA_m_, and H3 HA_m_ (From the top). The results of the final SEC and SDS-PAGE, SEC-MALS, and DSF of the HA mutant monomers are shown (From the left). Their molecular masses agreed with the value of 70.1 kDa, 62.5 kDa, and 85.0 kDa, respectively, taking into account glycosylation. The flow rates in SEC and SEC-MALS were 0.4 ml/min. Differential scanning fluorimetry transition curves are shown for the HA trimers (in blue) and mutant monomers (in purple). In case of H1 HA_m_, the recombinant H1 HA derived from CA04 is a monomer, whose T_m_ is lower than the H1 HA_m_.

All three PCNA subunits, PCNA1, PCNA2 and PCNA3, have a similar structure and length (∼250 amino acids), with a sequence similarity of 44−45%, and self-assemble into PCNA heterotrimers with affinities in the sub-μM to sub-nM range^27,29^. The mutant HA monomers, H1 HA_m_, B HA_m_ and H3 HA_m_, were fused to the scaffold subunits at the N-terminus, via SGG linker, to produce H1 HA_m_-PCNA1, B HA_m_-PCNA2, and H3 HA_m_-PCNA3, respectively (Fig. 1A & Supplementary Fig. S1). The sequences of all HA_m_-PCNA expression vectors were confirmed (Supplementary Table 1 & 2). Each assembly component was expressed in insect SF9 cells or mammalian HEK 293F cells and purified using His-tag affinity chromatography, ion exchange chromatography, and SEC (Supplementary Fig. S3A). The purified components were then mixed sequentially to form mosaic multivalent nanoparticles, 3HA-np. Self-assembly readily occurred irrespective of the cell system that was used for production (Supplementary Fig. S3B).

### Uniform mosaic display of antigens

Purified H1 HA_m_-PCNA1, B HA_m_-PCNA2, H3 HA_m_-PCNA3, and 3HA-np reacted with anti-H1, -B, and -H3 HA mAbs, as expected, suggesting uniform co-presentation of H1 HA_m_, B HA_m_, and H3 HA_m_ on 3HA-np (Fig. 2A & Supplementary Fig. S4). The SEC-MALS results showed that 3HA-np has molecular weights of 267.0 kDa (±12.1%) and 246.1 kDa (±9.9%) from mammalian and insect cells, respectively (Fig. 2B). H1 HA_m_-PCNA1, B HA_m_-PCNA2, and H3 HA_m_-PCNA3 had molecular weights of 91.0–103.5 kDa and 82.0–96.1 kDa from mammalian and insect cells, respectively (Supplementary Fig. S5). The mosaic 3HA-nps were visualized with atomic force microscopy (AFM) and transmission electron microscopy (TEM) and have an estimated overall size of 50-60 nm with three protruding HA antigens (Fig. 2C). Binding affinity data of 3HA-nps to sialic acid, a terminal sugar of N-glycans of fetuin, determined by biolayer interferometry (BLI) revealed dissociation constants (K_D_) of 1.4−1.5 × 10^-6^ M (Fig. 2D; Supplementary Fig. S6), whereas their affinity for asialofetuin was negligible (Data not shown). Binding of 3HA-nps to ConA, a mannose-binding lectin, was significantly reduced by treatment with Endo H, which was further reduced by the presence of mannan, a mannose-rich polysaccharide, suggesting that the 3HA-nps produced in insect or mammalian cells are N-glycosylated (Supplementary Fig. S7). Overall, the mammalian 3HA-np showed slightly higher affinity to sialic acid than the insect one, with lower binding to ConA, which nevertheless suggests that the 3HA-nps contained high levels of mannose glycans.

**FIGURE 2.**
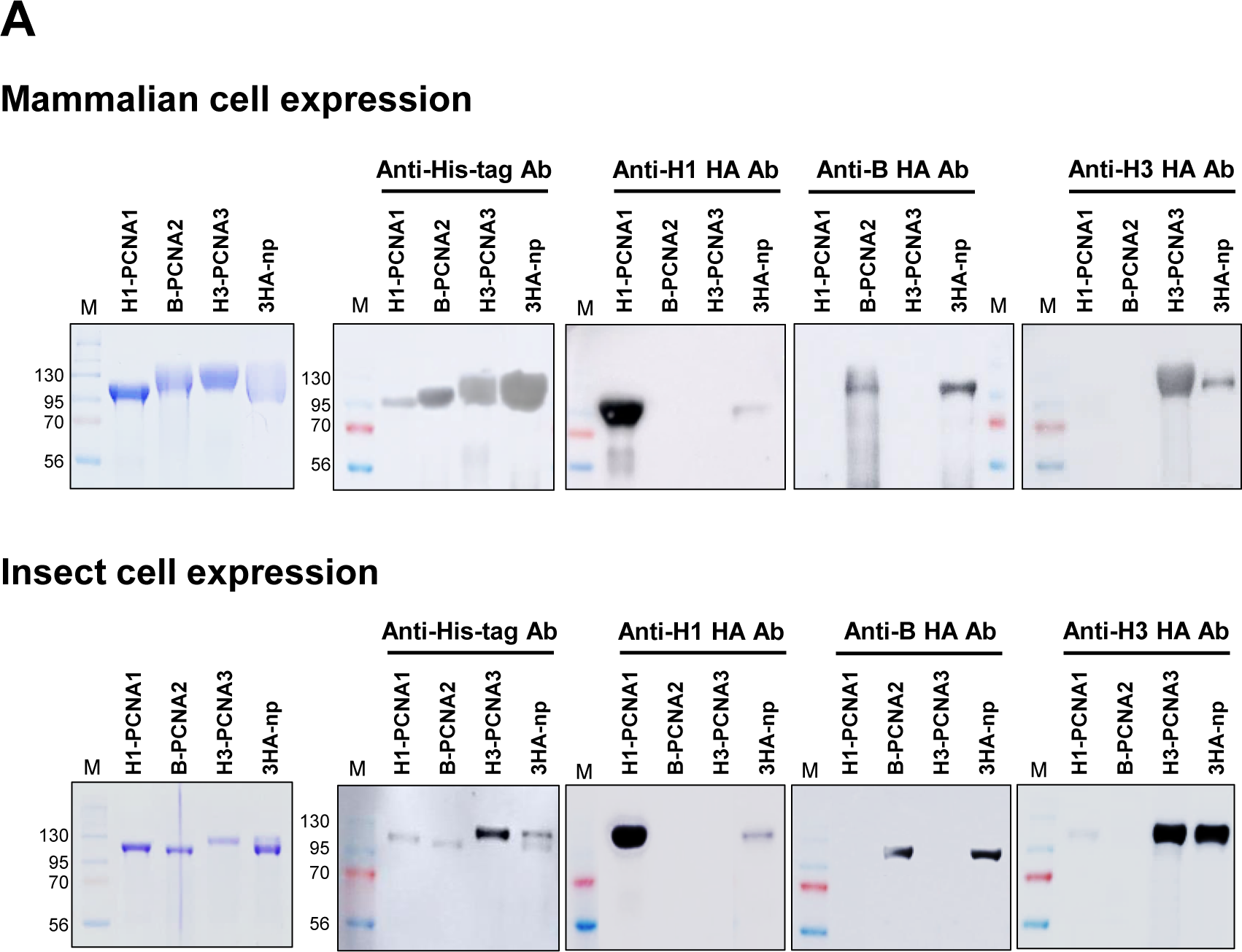
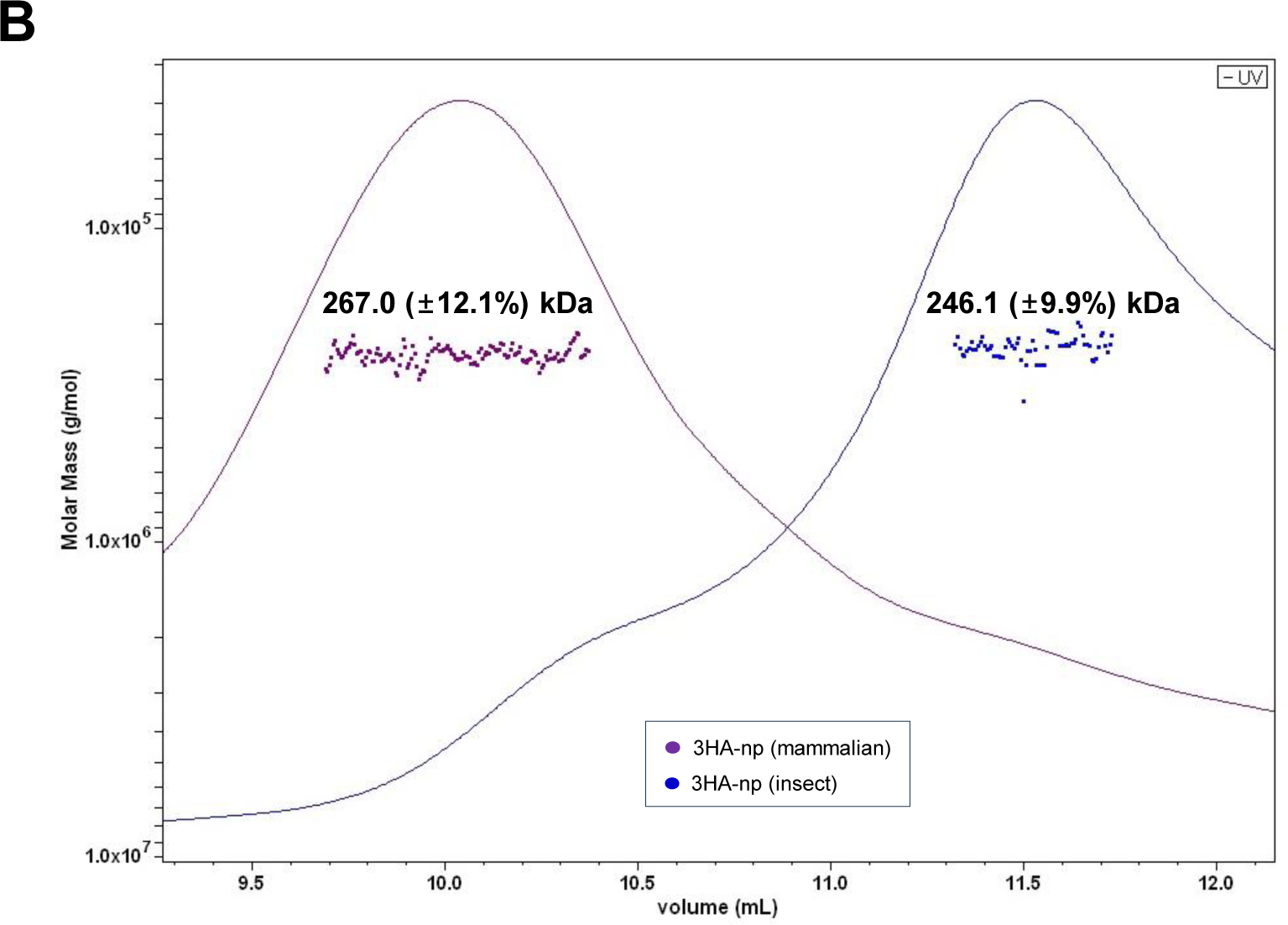
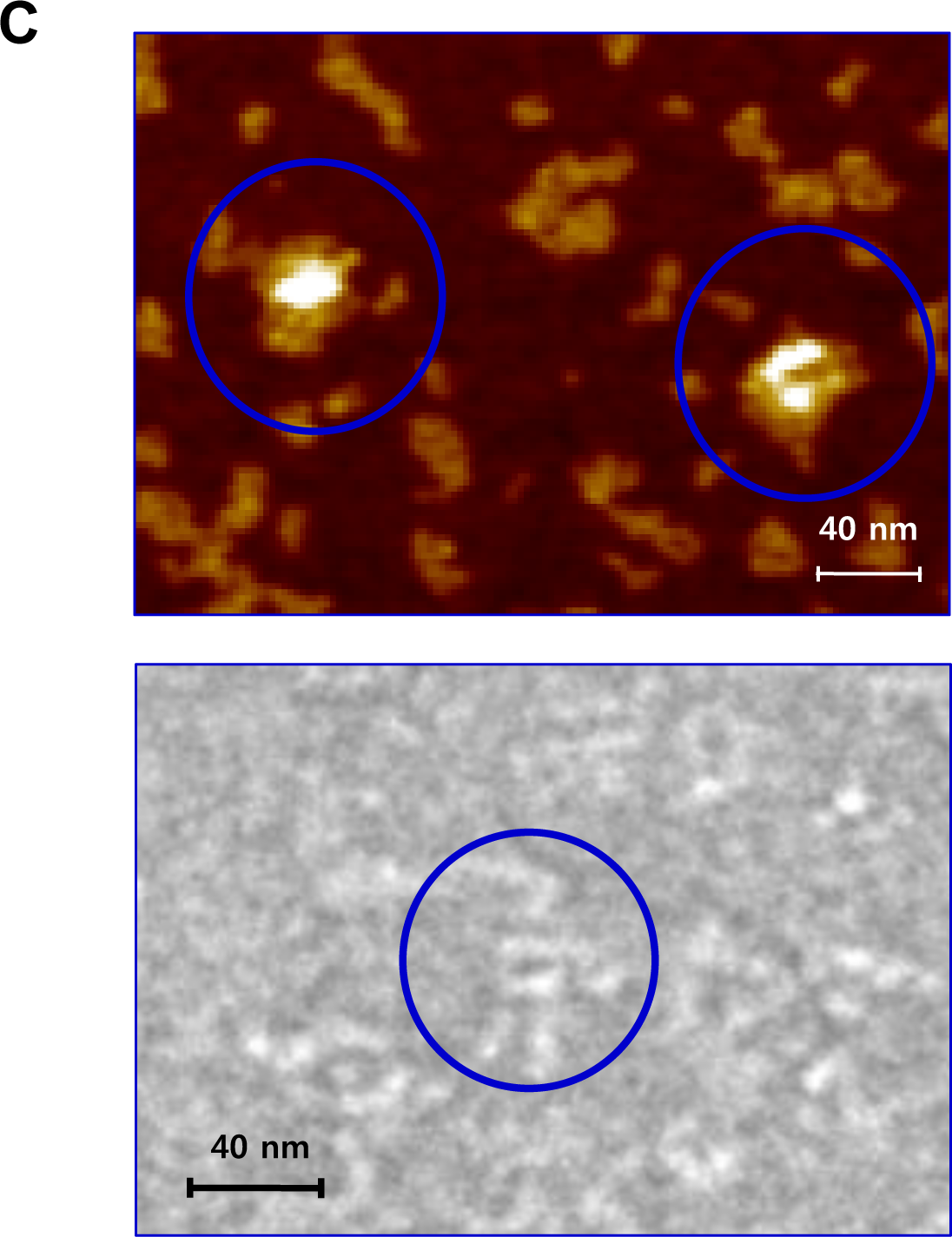
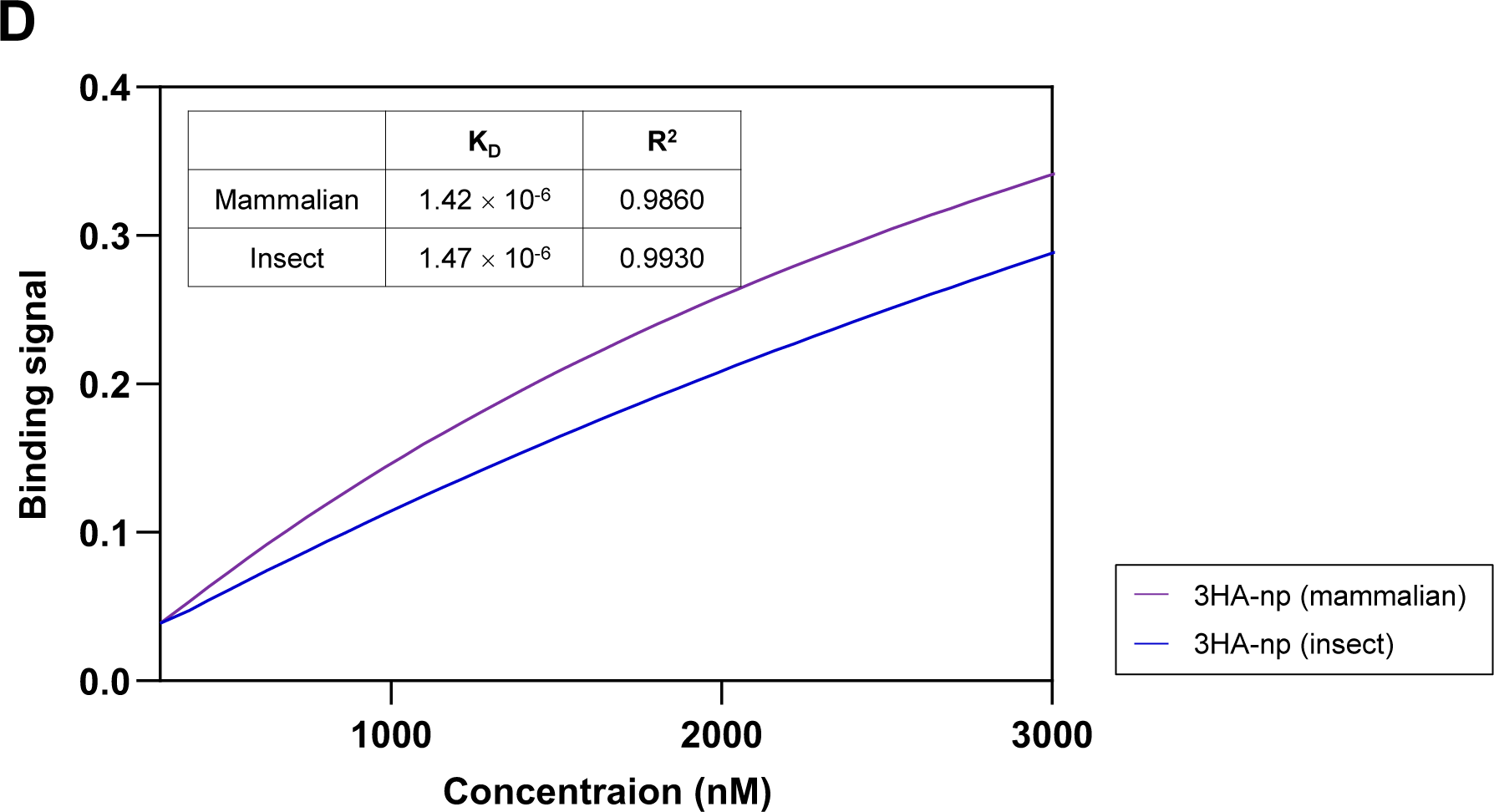
Antigenic and biophysical characterization of 3HA-np. (A) SDS-PAGE and Western blot assay results of the purified components, H1 HA_m_-PCNA1, B HA_m_-PCNA2, H3 HA_m_-PCNA3, and 3HA-np, using anti-His-tag Ab, anti-H1 HA Ab, anti-B HA Ab, and anti-H3 HA Ab from the mammalian (upper panel) and insect (lower panel) cell expressions. (B) Purified 3HA-np was characterized using SEC-MALS to show the molecular weights of 267.0 (±12.1%) and 246.1 (±9.9%) kDa from the mammalian and insect cell expressions, respectively. (C) AFM and TEM images of the mammalian 3HA-np with scale bars of 40 nm. (D) Determination of binding affinities of 3HA-nps to fetuin using BLI. Binding to fetuin was performed using Ni-NTA biosensors immobilized with the 3HA-np.

### Stability of 3HA-np

The stability of the components, H1 HA_m_-PCNA1, B HA_m_-PCNA2, and H3 HA_m_-PCNA3 and the assembled 3HA-np was monitored at −80℃, 4℃, 25℃, and 37℃ for 28 days. They were resistant to degradation at −80℃, and incubation of 3HA-np at 4℃ showed excellent stability, also suggesting that the mammalian cell produced fusion proteins showed slightly better stability (Fig. 3 and Supplementary Fig. S8). In contrast, severe degradation of 3HA-np was observed at 25℃ and 37℃ after 1 day. The stability of 3HA-np was affected by the least stable components upon assembly that are particularly B HA_m_-PCNA2 and H3 HA_m_-PCNA3. The mammalian 3HA-np remains as stable as the least stable components, showing a stability at 4℃ for at least a week.

**FIGURE 3.**
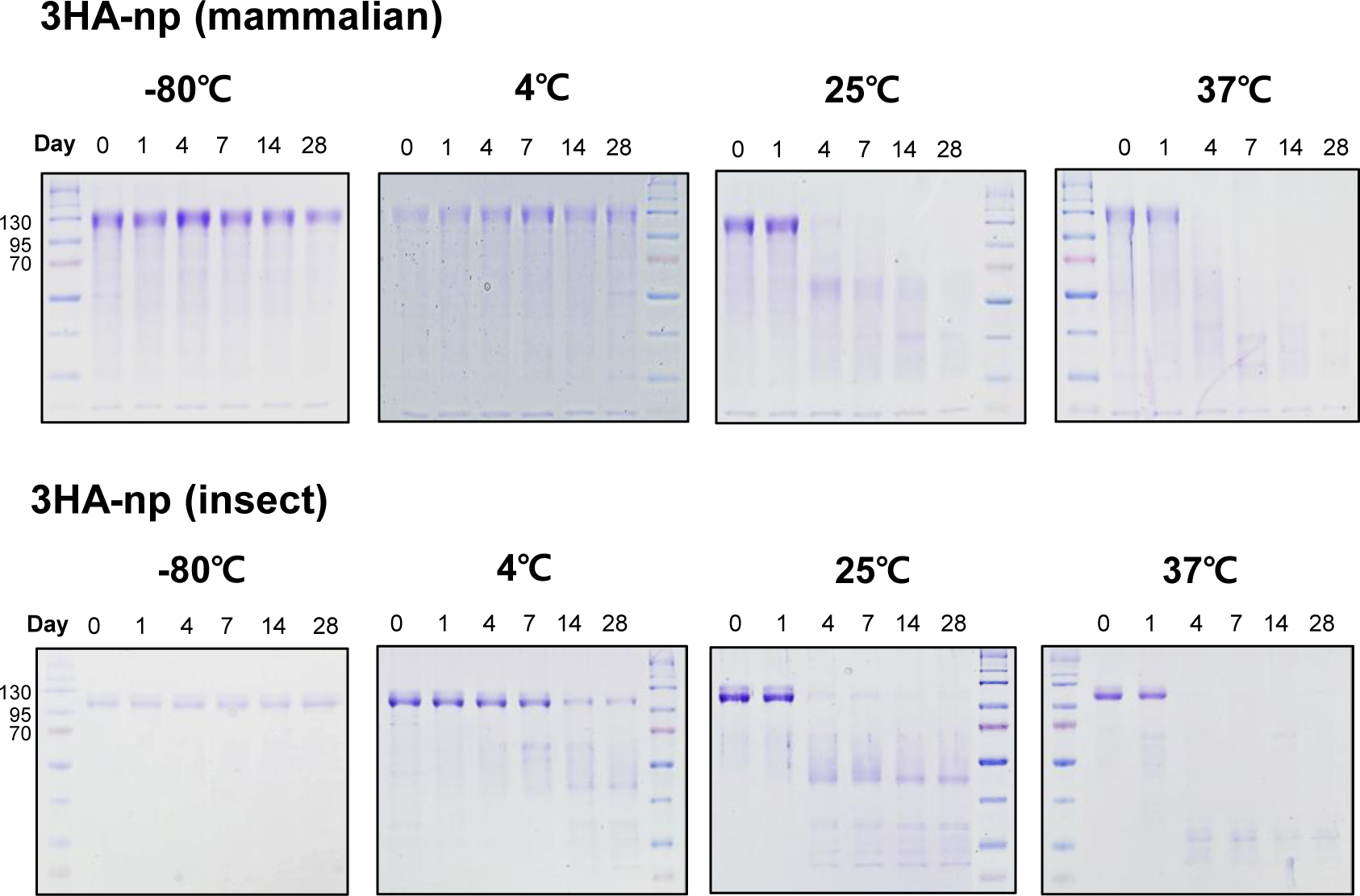
Time-course stability of 3HA-np. Stability of purified 3HA-nps expressed in mammalian (upper panel) and insect (lower panel) cells. SDS-PAGE results were monitored at −80℃, 4℃, 25℃, and 37℃ over a 28-day storage period (0, 1, 4, 7, 14, and 28 days).

### 3HA-np induces broad immune responses against influenza challenges in mice

Next, BALB/c mice were immunized with a mixture of HA monomer antigens (H1 HA_m_-PCNA1, H3 HA_m_-PCNA3 and B HA_m_-PCNA2), 3HA-np derived from insect cells, or 3HA-np derived from mammalian cells, by an intramuscular injection in the presence of AddaVax adjuvant in a prime-boost regimen (Fig. 4A). Three weeks after the boost immunization, the mice were challenged with 5LD_50_ of X47 (H3N2) or A/Puerto Rico/8/1934 (H1N1) (PR8) virus. All challenged mice experienced comparable body weight loss. Whereas mice that had been immunized with the HA mixture were only partially protected, all 3HA-np immunized mice survived the challenge with X47 or PR8 (Fig. 4A).

**FIGURE 4.**
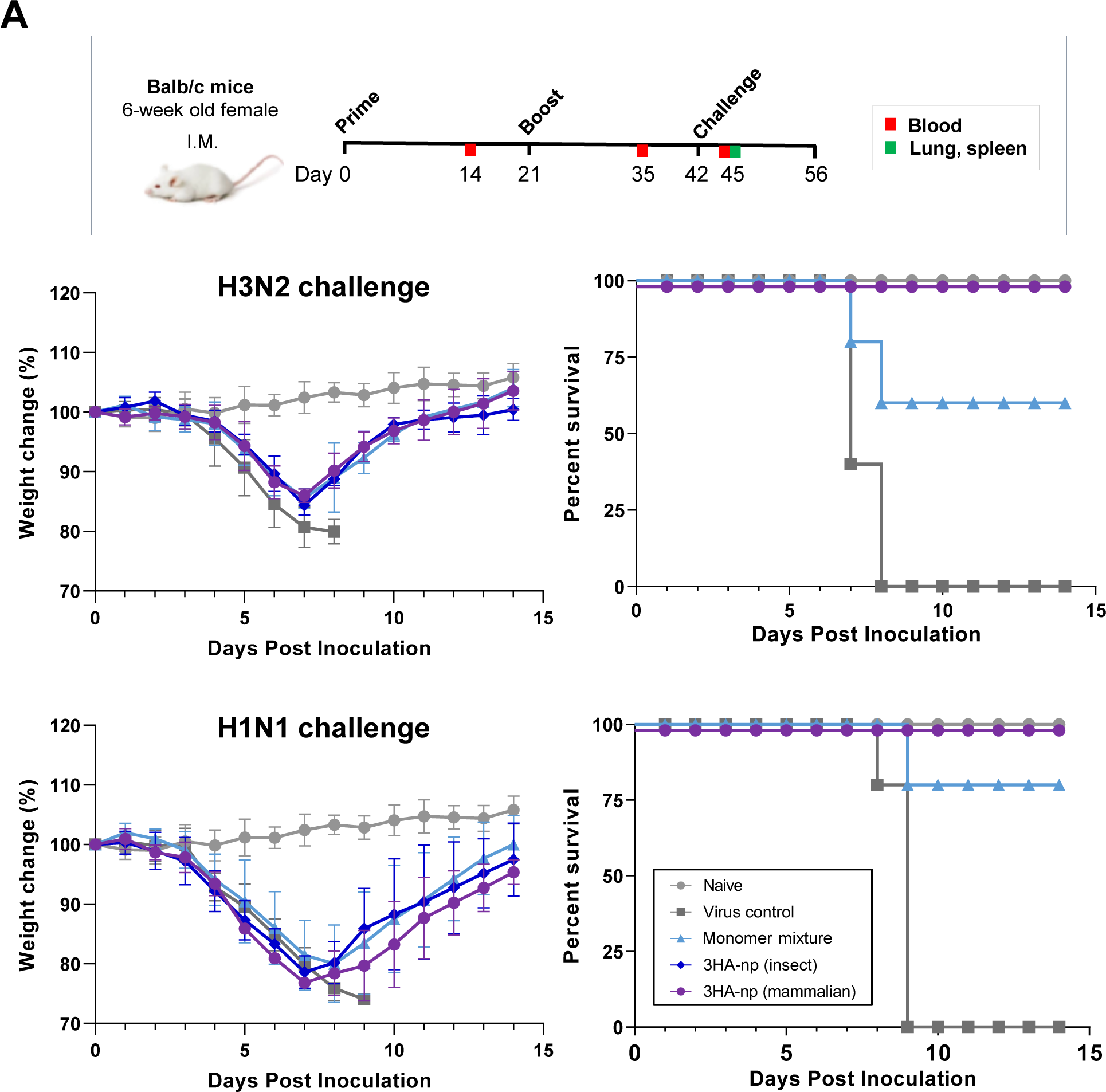
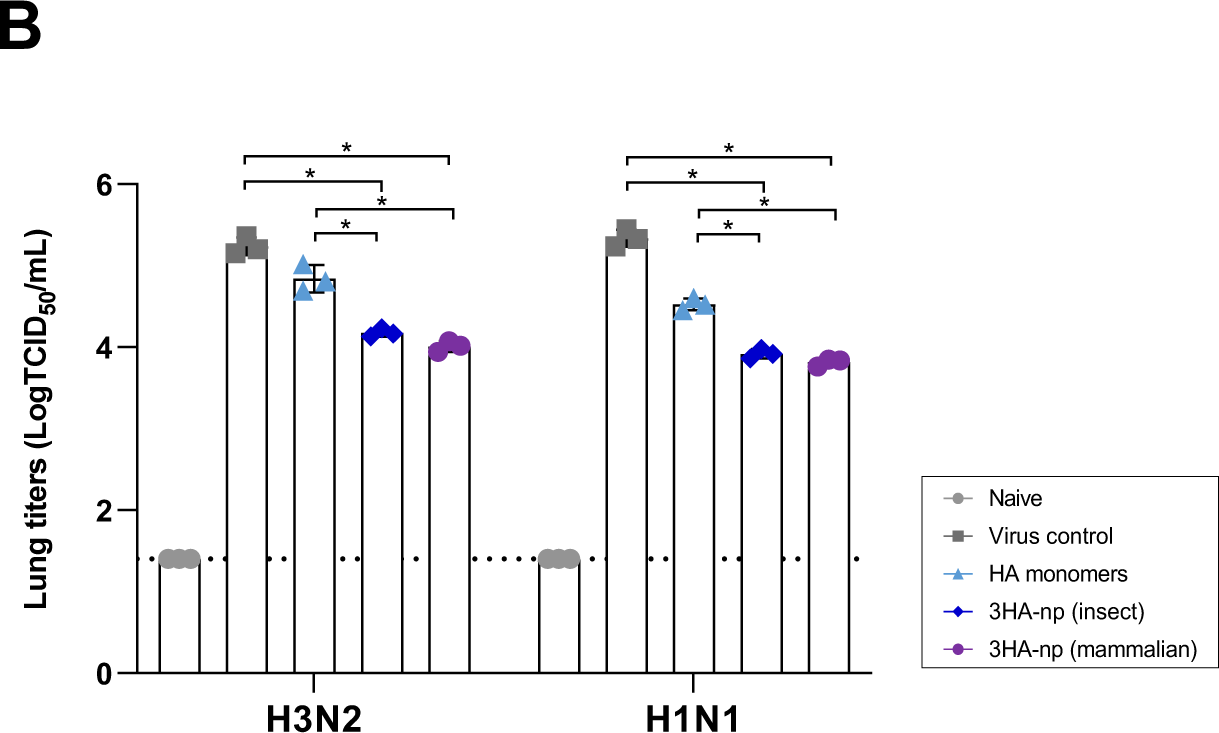
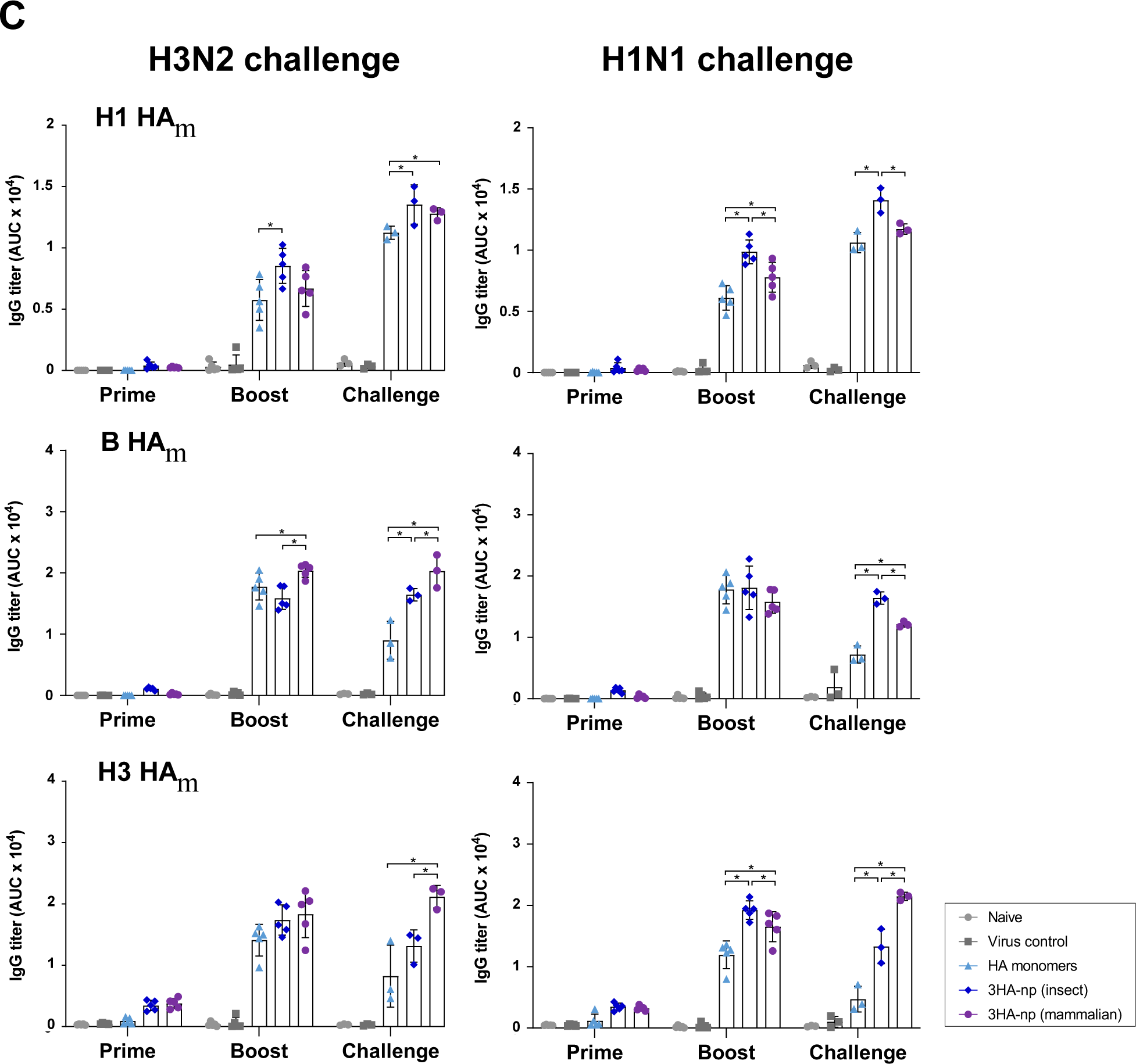
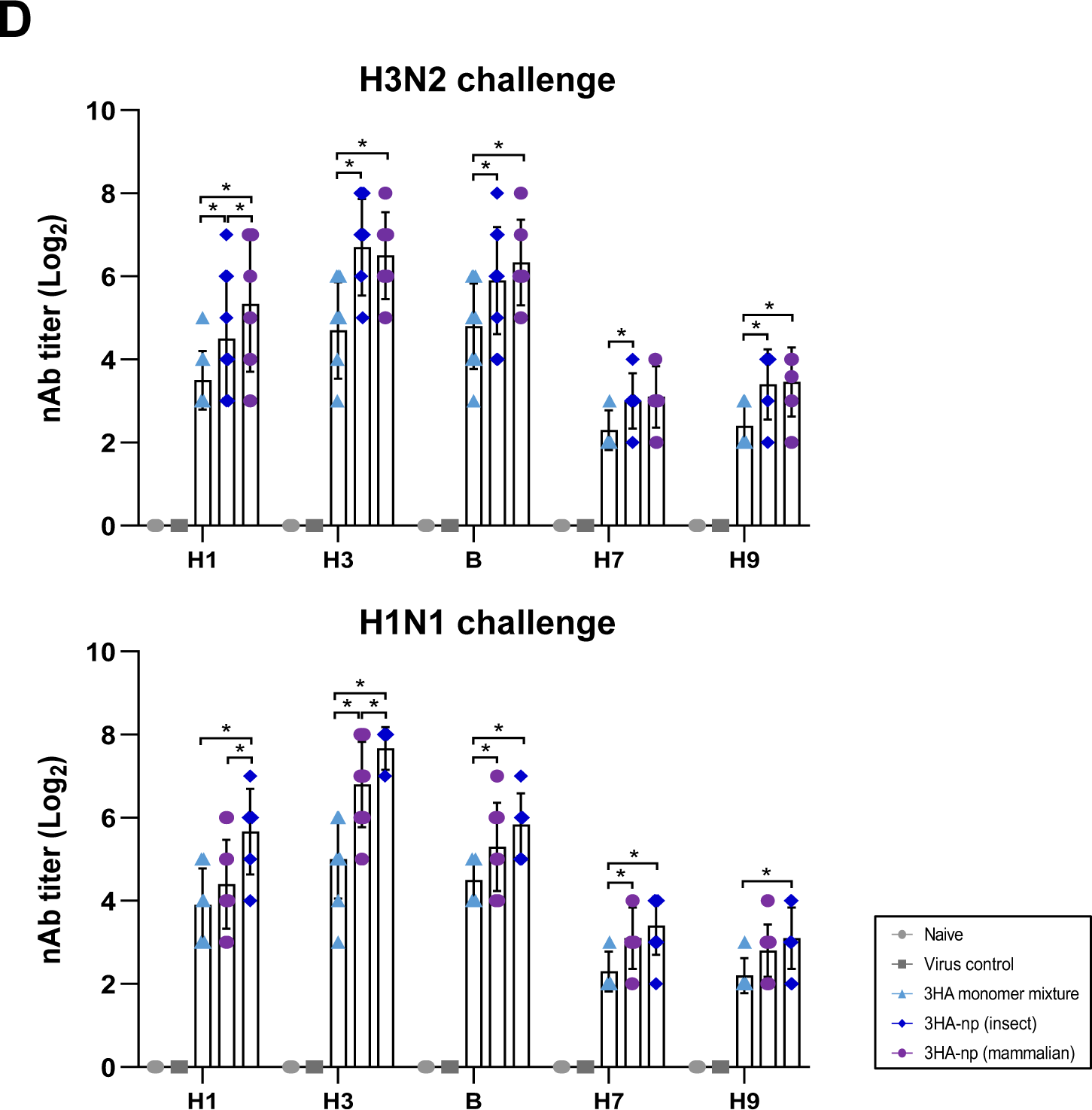
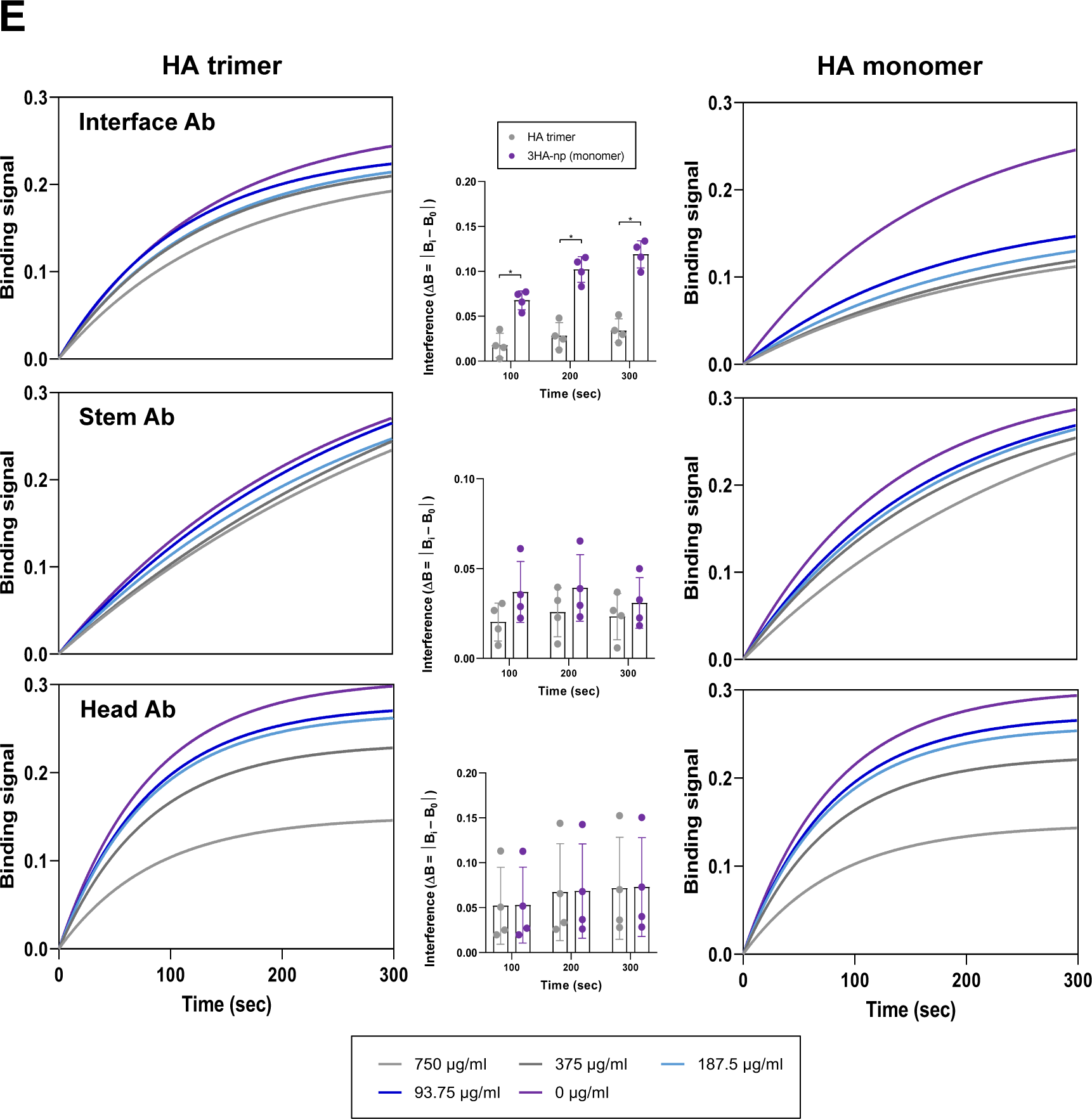
Protective efficacy and immunogenicity of 3HA-np against H3N2 and H1N1 challenges in mice. (A) Design of a challenge study with BALB/c mice (upper panel). The mice were immunized intramuscularly with 3 μg for prime and 5 μg for boost of antigens in five groups: the naïve, virus control, a mixture of HA monomer antigens (H1 HA_m_, H3 HA_m_ and B HA_m_), and 3HA-nps from mammalian and insect cells. After 3 weeks of boost immunization, mice were intranasally inoculated with A/reassortant/X-47 (H3N2; X47) or A/Puerto Rico/8/1934 (H1N1; PR8) virus. Weight change and percent survival of mice challenged with H3N2 (middle panel) and H1N1 (lower panel) were monitored up to 14 dpi. (B) Viral titers in mice lungs collected from 3 mice per group following the H3N2 or H1N1 challenge. The titers were measured using plaque assays. (C) Anti-HA Ab titers in immunized mice sera after prime, boost, and challenges with the H3N2 or H1N1 virus, using ELISA assays. The antigens, H1 HA_m_-PCNA1, B HA_m_-PCNA2, and H3 HA_m_-PCNA3, were incubated with serially diluted sera, and serum IgG responses are shown in AUC at 450 nm. (D) NAb titers in mice sera after challenges with H3N2 (upper panel) or H1N1 (lower panel) measured by microneutralization (MN) assay. Influenza viruses, A/Korea/01/2009 (H1N1), B/Florida/04/2006, A/Brisbane/10/2007 (H3N2), A/chicken/Korea/L433/2018 (H9N2), and A/duck/Korea/557/2016 (H7N7) were used. (E) Competition assays for Abs in mice sera with mAbs, GC1517, CR9114, and D2 H1-1/H3-1. The H3 HA wild type trimer (left panel) or monomer (right panel) was immobilized to the Ni-NTA biosensor. MAbs, GC1517, CR9114, and D2 H1-1/H3-1, directed against the head, stem, and interface epitopes, respectively, were used to identify their competitive effects on Abs in mice sera in the range of 0 – 750 μg/mL. Interference differences in binding responses are represented by ΔB = |B_i_ - B_0_|, where B_i_ and B_0_ are the concentrations at 93.75−750 μg/mL and 0 μg/mL, respectively (center panel). The mean values are indicated by a bar, where each value from individual sera is shown as a symbol. The data were compared using the Mann-Whitney U test (*p < 0.05), and statistically significant differences were indicated only for competition with the interface Ab.

The 3HA-np groups showed lower viral titers ranging from 1.1 to 1.3 log TCID_50_/mL in the lung tissue collected at 3 days post-infection (dpi) for both the H3N2 and H1N1 challenges, compared to the virus control and the HA monomer mixture group (Fig. 4B). The viral titers were only partially inhibited in the mixture group. Enzyme-linked immunosorbent assay (ELISA) was performed with individual sera, using purified H1 HA_m_-PCNA1, B HA_m_-PCNA2, and H3 HA_m_-PCNA3 as coated antigens. The groups that had been immunized with either mammalian or insect cell-derived 3HA-np had significantly higher antigen-specific Ab titers than the HA monomer mixture group, except for the B HA monomer titers (Fig. 4C).

We then evaluated the bnAbs in the post-challenge sera for microneutralization (MN) activity against a broad range of influenza virus strains, including A/Korea/01/2009 (H1N1), A/Brisbane/10/2007 (H3N2), B/Florida/04/2006, A/duck/Korea/557/2016 (H7N7), and A/chicken/Korea/L433/2018 (H9N2). The immune sera from mice in the 3HA-np groups showed significantly higher neutralizing potency than those from the monomer mixture group, regardless of the challenges (Fig. 4D). Importantly, they revealed higher level of nAb titers than the monomer mixture group, suggesting an improved neutralization breadth exhibited by 3HA-np against diverse influenza virus HA subtypes, including H7 and H9 viruses. To test whether binding to the conserved stem or interface epitopes correlated with increased cross-reactivity, competition assays were performed using BLI to characterize Ab repertoires in mice sera. MAbs, GC1517, CR9114, and D2 H1-1/H3-1, that are directed against spatially distinct epitopes, the head, stem, and interface, were used as competitive binders, respectively^18,23,31,32^. D2 H1-1/H3-1, specific to the monomer-monomer interface epitope, showed the most significant competition to the 3HA-np sera among the groups, suggesting that significantly higher titers of interface-specific Abs were induced in the sera of 3HA-np immunized mice, due to the protruding HA monomers on 3HA-np (Fig. 4E and Supplementary Fig. S9). The competition was more pronounced in H3 HA than H1 HA immobilized. The stem Ab, CR9114, also seemed to have a competition to the 3HA-np sera, whereas there was little competition by the head-specific Ab. GC1517 was known to neutralize a broad range of H1 subtype influenza A viruses^32^, which was unexpectedly bound to H3 HA, albeit weaker than H1 HA (Supplementary Fig. S10). Taken together, our results demonstrated that immunization with 3HA-np provided 100% protection against lethal challenges of both H1N1 and H3N2 viruses with consistently high nAb levels, and strongly suggest that the 3HA-np markedly elicits crAb responses, including interface-specific Abs.

Splenocytes from the immunized mice stimulated with a peptide pool of HA from the H1N1 influenza virus A/California/04/09 were used for enzyme-linked immunospot (ELISpot) assay. The number of interferon (IFN)-γ producing T cells was significantly higher in the 3HA-np groups than the HA monomer mixture group (Supplementary Fig. S11). The results further support that the mosaic multivalent antigen 3HA-np induced significantly high IFN-γ producing T cell responses against the H1 and H3 challenges, irrespective of the eukaryotic cell expression system used.

### 3HA-np elicits significantly high nAb titers in ferrets against sequential heterologous virus challenges

To further evaluate the breadth of immunogenicity and protective efficacy, four-month-old ferrets were divided into five groups of native, virus control, H3 HA monomer, and insect and mammalian 3HA-nps. The ferrets were given intramuscular prime-boost immunizations with a 3-week interval (20 μg and 25 μg antigens with Addavax, respectively) (Fig. 5A). At two weeks post-boost, groups of immunized ferrets were challenged intranasally with H3N2 (A/Perth/16/2009) followed by H1N1 (A/California/04/2009) virus challenges two weeks later. When monitored for body weight and body temperature for 8 dpi, the ferrets immunized by 3HA-np exhibited a significant increase of body weight of 10-20% and 5-10% upon sequential infections of H3N2 and H1N1 viruses, respectively (Fig. 5A). By contrast, the naïve and virus control ferrets showed little change or even a slight reduction in body weight. Immunization with the 3HA-nps effectively prevented an increase in body temperature after H3N2 and H1N1 challenges, compared to the significant increase in the H3 HA monomer and virus control groups. The viral titers in nasal wash, collected at 1, 3, and 5 dpi upon virus challenges, showed little difference among the groups at 1 dpi (Fig. 5B). At 3−5 dpi, however, ferrets in the 3HA-np groups had significantly lower virus titers in the H3N2 and H1N1 challenges than other groups.

**FIGURE 5.**
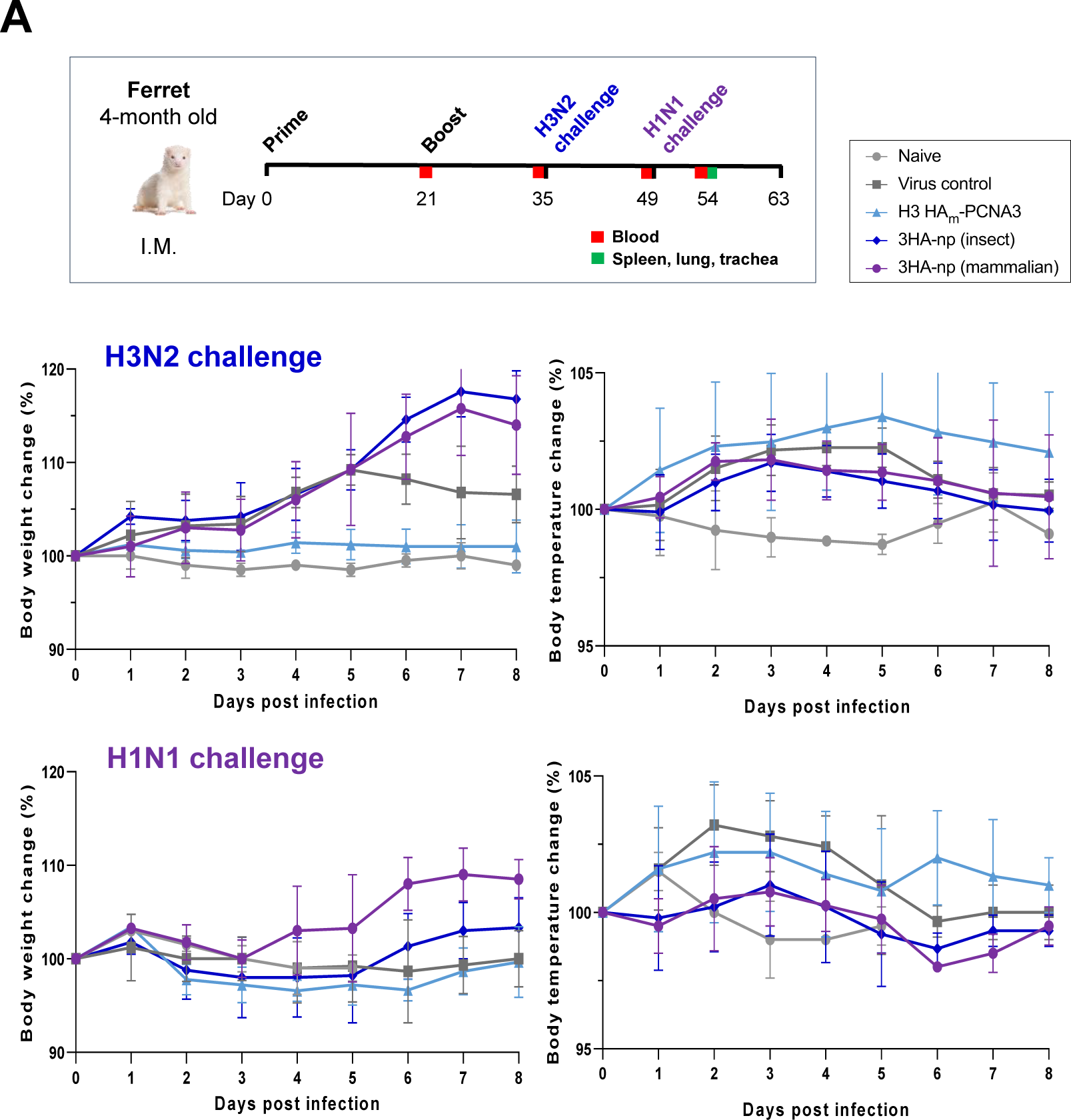
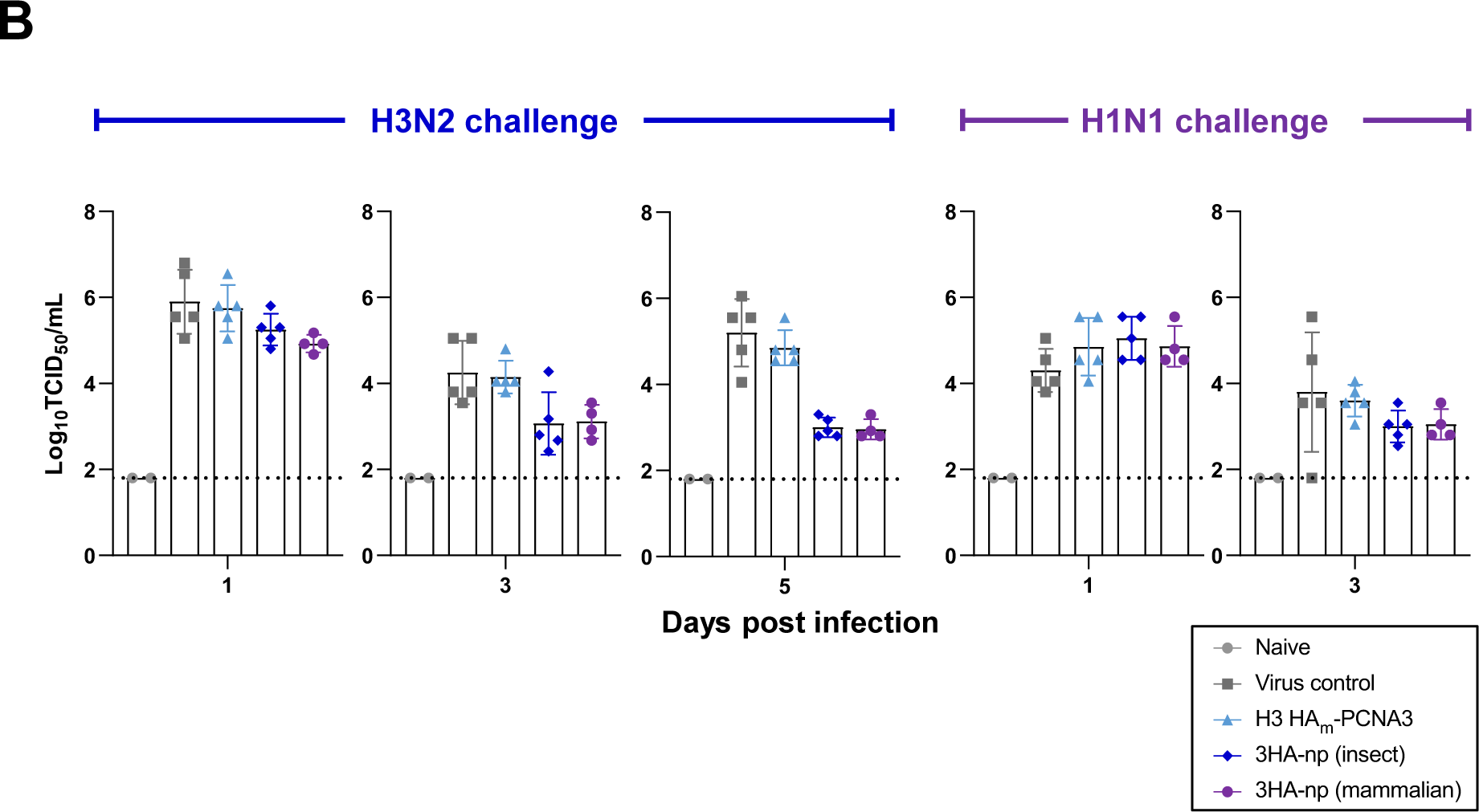
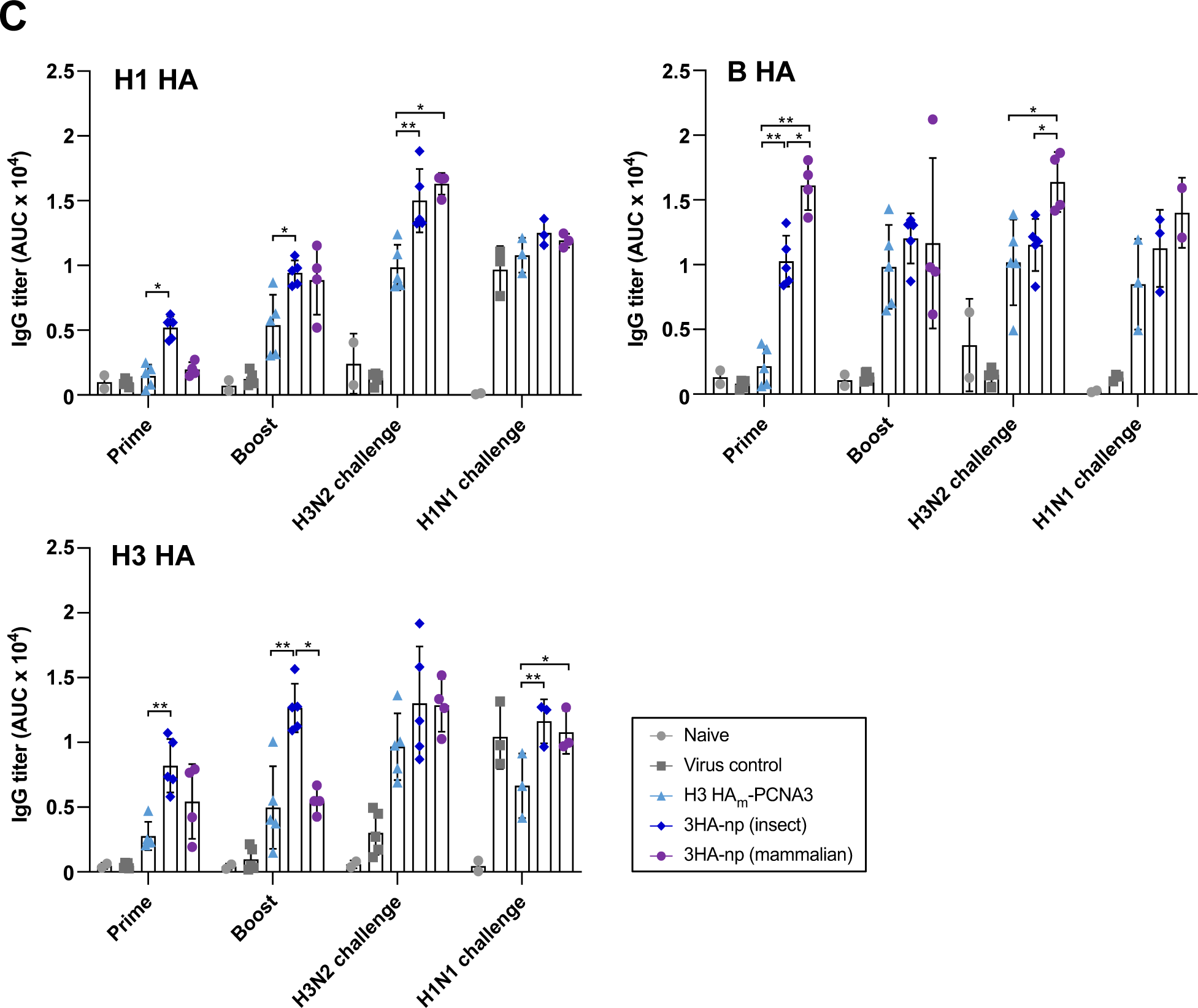
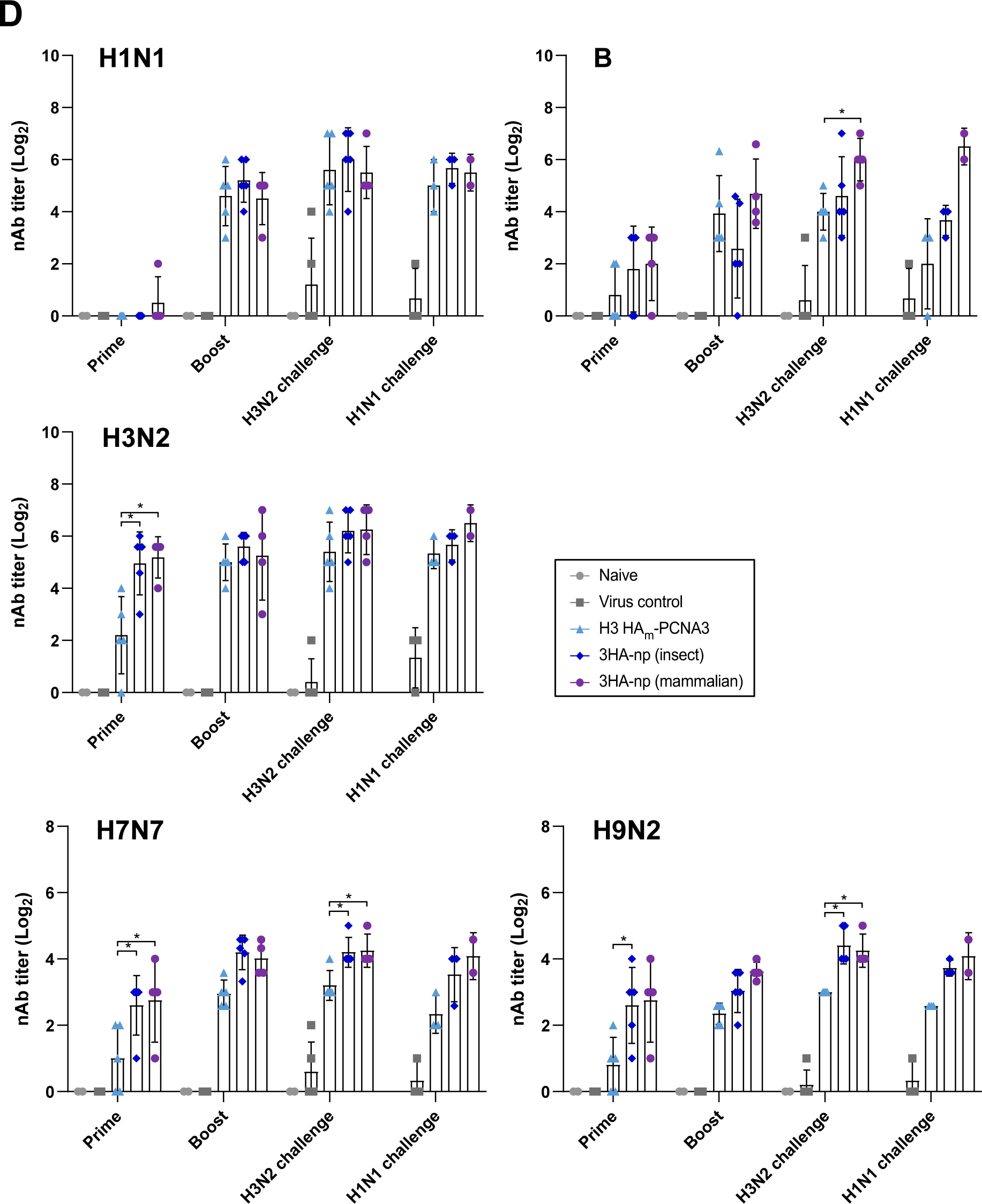
Protection and immunogenicity of 3HA-np in ferrets. (A) Design of prime-boost immunization and challenge studies in ferrets (upper panel). Four-month-old ferrets were given two intramuscular immunizations with a 3-week interval (20 μg and 25 μg antigens with Addavax, respectively). They were divided into five groups of the naïve, virus controls, H3 HA monomer from mammalian 293F cell, and 3HA-nps from mammalian and insect cells. At two weeks post-boost, all groups except for the naïve group were challenged with serial intranasal inoculation (6.0 Log TCID_50_/ml) of heterologous influenza H3N2 (A/Perth/16/2009) and mouse-adapted H1N1 (A/California/04/2009) viruses with a two-week interval. The body weight and temperature of the ferret were monitored up to 8 dpi after the H3N2 (middle panel) or H1N1 (lower panel) challenge. Ferret sera were collected at 3 weeks post-prime, 2 weeks post-boost, 2 weeks post-H3N2 challenge, and 5 days post-H1N1 challenge. (B) Viral titers in nasal wash in ferrets. Nasal wash samples were collected at 0, 1, 3, and 5 dpi after H3N2 (left panel) or H1N1 (right panel) challenge. MDCK cells were infected with 50 μL of the 10-fold serially diluted supernatant of the nasal wash for 1 hr. Virus-induced cytopathic effects were measured, and viral titers were expressed in log TCID_50_/mL. Viral titers in the nasal wash were detected below the detection limit after 7 or 5 dpi in H3N2 and H1N1 challenges, respectively. (C) Antigen-specific Ab titers in ferret sera determined by ELISA assays. The antigens, H1 HA_m_-PCNA1, B HA_m_-PCNA2, and H3 HA_m_-PCNA3, were incubated with serially diluted sera, and serum IgG responses are shown in AUC at 450 nm. (D) NAb titers were measured by microneutralization assay. A/Korea/01/2009 (H1N1), B/Florida/04/2006, A/Brisbane/10/2007 (H3N2), A/chicken/Korea/L433/2018 (H9N2), and A/duck/Korea/557/2016 (H7N7) influenza viruses at 2 log10 TCID_50_/mL were mixed with diluted serum at a 1:1 ratio prior to 1 hr incubation at 37°C. Mean nAb titers were calculated and indicated by a bar, where each value from individual ferret sera is presented as a symbol, and the standard deviation is indicated by a vertical line. The data were compared using the Mann-Whitney U test (*p < 0.05; **p < 0.01). Statistically significant differences were indicated only among different immunogen groups.

Using H1 HA_m_-PCNA1, B HA_m_-PCNA2, and H3 HA_m_-PCNA3 as coated antigens, ELISA performed with the post-prime, post-boost and post-challenge sera indicated that ferrets immunized with 3HA-nps had significantly higher antigen-specific Ab titers, irrespective of H1N1 and H3N2 virus challenges (Fig. 5C). We next assessed nAb titers against authentic influenza virus strains, A/Korea/01/2009 (H1N1), A/Brisbane/10/2007 (H3N2), B/Florida/04/2006, A/duck/Korea/557/2016 (H7N7), and A/chicken/Korea/L433/2018 (H9N2). Consistent with the nAb titers in mice, 3HA-np elicited high neutralizing potency and breadth against diverse influenza virus isolates, including H7N7 and H9N2 strains (Fig. 5D). Furthermore, the number of IFN-γ producing T cells in ferret splenocytes determined by ELISpot assays was more than 2-fold higher in the mammalian 3HA-np group than the H3 HA monomer group (Supplementary Fig. S12). Taken together, both mice and ferrets immunized by 3HA-np demonstrated broad immunogenicity and protective potential against H1N1 and H3N2 influenza viruses.

## DISCUSSION

A pool of broadly crAbs against influenza viruses have been shown to target discrete antigenic epitopes of HA with high conservation across multiple subtypes: the receptor binding site, the occluded intermonomer interface, and the stem regions of hydrophobic groove, fusion peptide and membrane-proximal anchor^33,34^. The rational design of conserved epitopes is technically challenging, while a panel of headless, chimeric, hyperglycosylated, scaffold-based, and virus-like particle-based antigens has been explored, primarily focusing on highly conserved stem epitopes^14,15,20,21,35−37^. Sequential vaccination with chimeric HAs, consisting of head domains of different subtypes combined with the same subtype stem, underwent a successful phase 1 clinical trial^16^. Further, co-display of heterotypic antigens by designed scaffold or Spy-tag-mediated fusion chemistry elicited avidity-mediated enhancement of broad Ab responses to target conserved epitopes^25,26^. In this study, we introduce a novel approach to display heterotypic H1, H3 and B HA monomers, instead of trimers, and to incorporate heterotypic antigens into self-assembled scaffolds. Juxtaposition of the heterotypic antigens in the structure of 3HA-np, appeared as a ring-shaped disk with three protruding HA monomers, was shown to provide protection against mortality for both H1N1 and H3N2 virus challenges in a mouse model and to markedly elicit broader cross-reactive immune responses in mice and ferrets against H7 and H9. It was previously reported that mosaic nanoparticles based on SpyCatcher-based platforms, presenting heterotypic conjugation of 8 different HA trimers, elicited strong Ab responses, but did not consistently elicit broader immune responses than the admixture of homotypic particles^38^. Given that no other methods currently ensure mixing uniformity of heterotypic antigens on a single nanoparticle, it is likely that the mosaic nanoparticle could not clearly present uniform distribution of 8 heterotypic antigens and consequently elicit immune responses of greater breadth. In contrast, our mosaic nanoparticle platform based on self-assembly of heterotrimeric subunit scaffolds proved to achieve a uniform distribution of heterotypic antigens, which potentially increases the breadth and magnitude of the immune responses.

Seasonal influenza vaccines are mainly produced in eggs and expressed in cell-based systems such as insect and mammalian cells. They differ in glycosylation profiles and sulfation diversity in N-glycans, which may influence immunological properties^39^. When immunogenicity of recombinant HA antigens was tested in chickens and mice, mammalian-expressed HAs induced significantly higher nAb titers than insect cell HAs. However, the glycosylation difference did not affect the breadth of nAb responses^40^. In this study, the epitopes of monomeric HA antigens on 3HA-np were uniformly recognized by anti-HA mAbs, showing high potency and breadth of responses and even increased IFN-γ production in splenocyte derived from immunized mice, irrespective of mammalian and insect cells. More striking differences in Ab responses were observed between the groups of HA monomer mixture and 3HA-nps, rather than between the 3HA-nps produced in mammalian versus insect cells. Although the mammalian 3HA-np was found to be slightly superior to the insect cell produced 3HA-np in some of the tested anti-HA Ab and nAb responses, there were no significant and consistent difference between the two different cell expression systems.

The stability of HA is an important attribute of influenza viral fitness to the extent that HA can tolerate mutations for immune escape. The recombinant HA derived from A(H1N1) 2009 viruses revealed a monomeric nature and evolved to exhibit significantly more stable HA as they became seasonal strains circulating globally^41^. The mosaic 3HA-np signified an approach that allows the protruding monomeric HA antigens to expose the monomer-monomer interface occluded inside the HA trimer (Fig. 6). The interface epitope that is conserved across most subtypes of influenza A viruses was shown bound by the interface Abs^18^, which can provide a broad-spectrum protection against influenza viruses. Cross-competition assays using mice sera demonstrated that binding of D2 H1-1/H3-1 to the monomer-monomer interface epitope posed significant competition to the 3HA-np sera, which presumably reflects the increased exposure of the interface epitope by the stable protruding HA monomers on 3HA-np. Therefore, 3HA-np can induce strong and broad immune responses based on the mosaic and monomeric nature of heterotypic HAs. The mosaic multivalent nanoparticle is potentially able to provide cross-reactive protection against circulating and zoonotic influenza viruses. During the fusogenic process, the interface epitope is thought to be transiently exposed as HA undergoes a conformational transition, and a single-molecule Forster resonance energy transfer-imaging study reported that HA can exhibit reversible conformation changes at physiological pH that allow temporary interface exposure^18,42^.

**FIGURE 6.**
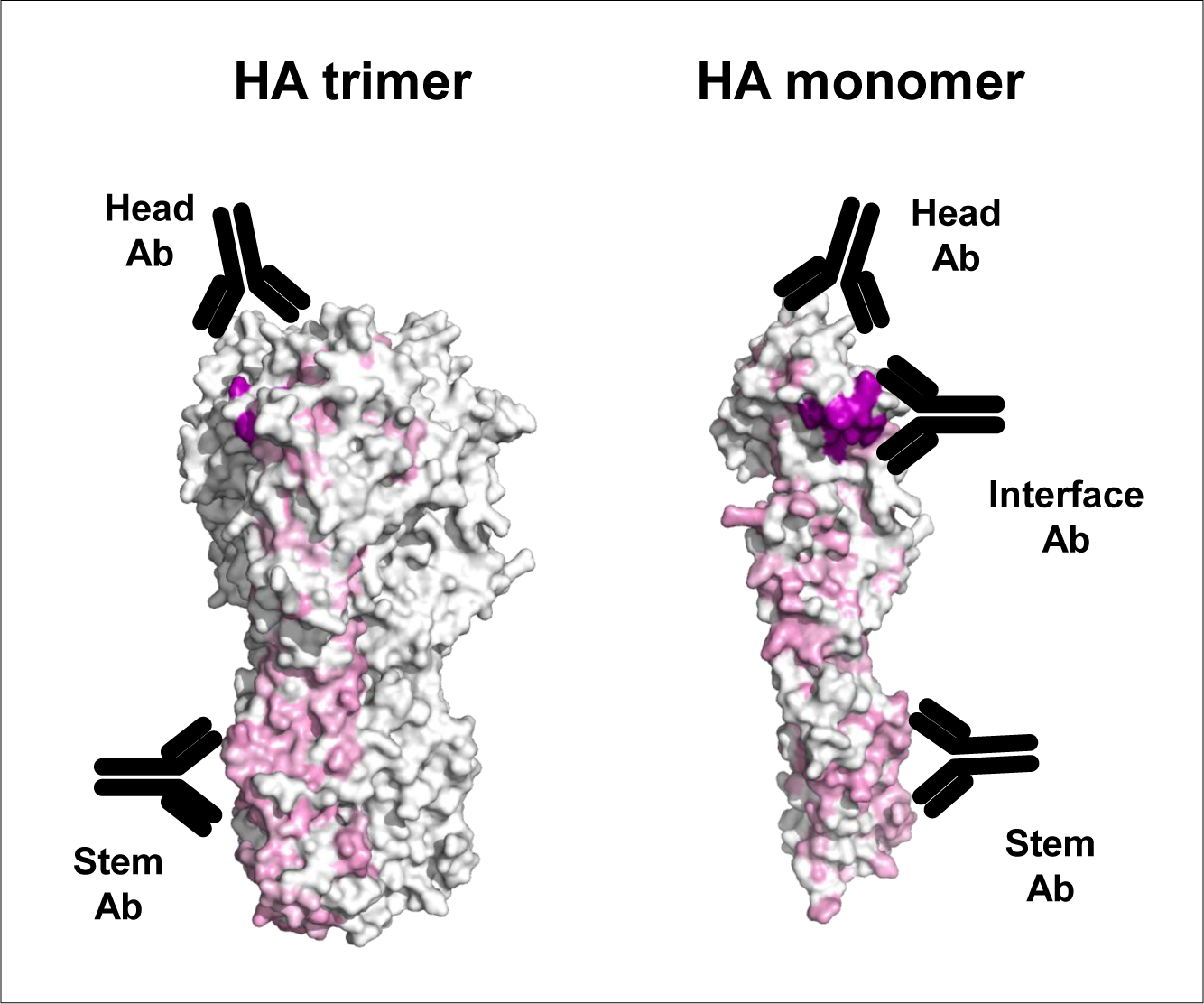
Overview of HA epitopes with conservation of amino acid residues. The HA trimer antigen present the head and stalk epitopes, whereas the HA monomer can additionally present occluded interface epitopes to which Abs bind, presented as black Y-shapes. Conserved amino acid sequence regions are highlighted in pink, mostly overlapped in the stem region, and the occluded interface epitope is highlighted in purple. The 3HA-np allows the protruding monomeric HA antigen to expose the monomer-monomer interface occluded inside the HA trimer. The HA models were generated from PDB 4WE8 with the amino acid sequence conservation calculated by ConSurf^44^.

Yearly updated influenza vaccines depend on a large amount of epidemiological, genetic, antigenic, and human serological data. Year-round surveillance and monitoring of influenza viruses are performed on a regular basis, together with collection and testing of circulating virus strains based on regional and global influenza laboratory network. Such a coordinated global effort leads to the selection of currently circulating virus strains to update seasonal influenza vaccines. Despite the arsenal of influenza virus seasonal vaccines, influenza morbidity and mortality remain high due to their variable and sometimes weak effectiveness against circulating viruses. Protection induced by seasonal vaccines against potential pandemic strains is sub-optimal, and broadly cross-reactive vaccines that can achieve a durable protection are therefore urgently needed. In this context, a universal influenza vaccine would be a game changer that abolish the need for seasonal vaccine updates and improve future pandemic preparedness. However, the development of universal influenza vaccines that induces broadly protective antibodies has been a challenge for many decades. In this study, the mosaic antigen platform offers an approach that allows the induction of cross-reactive B cell responses thereby providing an advantage over strain-specific homotypic antigens. We have particularly shown that the scaffold-based mosaic nanoparticle platform has the potential to provide a feature ensuring the uniform arrangement of three distinct antigens on nanoparticles.

## MATERIALS AND METHODS

### Design, expression, and purification of 3HA-np

For the 3HA proteins, the residues of A/California/04/2009 (H1N1) (CA04), B/Florida/4/2006 (FL04), A/Gyeongnam/684/2006 (H3N2) (Gy684) influenza viruses as well as PCNA1, PCNA2, PCNA3_S170V were amplified as described in Tables S1 & S2. For stable HA monomer (HA_m_) production, specific mutations were introduced into the H1, B, and H3 HA proteins (Fig 1A). Fusion proteins, H1 HA_m_-PCNA1, B HA_m_-PCNA2, and H3 HA_m_-PCNA3, were constructed with an SGG linker. They were expressed in mammalian Expi293F and insect Sf9 cells and purified by Ni-NTA affinity chromatography, ion-exchange chromatography, and SEC. The purified H1 HA_m_-PCNA1, B HA_m_-PCNA2, and H3 HA_m_-PCNA3 were sequentially mixed, and the assembled 3HA-np was further purified by SEC. Each HA-PCNA antigen was also constructed, expressed, purified, and used for assays.

### Characterization of antigens

Light scattering and refractive index were measured using WYATT-787-TS miniDAWN system. Negatively stained electron microscopic images were obtained under a Tecnai 20 transmission electron microscope operating at 120 kV, and AFM data were acquired using Multimode-Ⅷ at 0.3 Hz in a line. The binding of antigens to receptors were measured by bio-layer interferometry. Fusion proteins and 3HA-np were treated with Endo H for deglycosylation, and overall structures of HAs were superimposed using the Coot^43^. The amino acid sequence conservation was calculated at the ConSurf server^44^, based on multiple sequence alignment of 150 HA proteins, and conserved sites were presented on the structure of an H3 HA (PDB ID: 4WE8).

### Mouse immunization and challenge

BALB/c mice were divided into five groups of the naïve (n=5), virus control (n=10), a mixture of HA monomer antigens (H1 HA_m_, H3 HA_m_ and B HA_m_, n=10), and 3HA-nps derived from mammalian (n=8) and insect (n=10) cells and immunized in a prime-boost protocol with a three-week interval. Antigens were injected intramuscularly at a dose of 3 μg and 5 μg with AddaVax (InvivoGen) in 100 μL for prime and boost, respectively. After 3 weeks of boost immunization, mice were intranasally inoculated with 5 median lethal dose 50% of the influenza virus H3N2 (X47) or H1N1 (PR8), respectively, under general anesthesia. Throughout a monitoring period of 14 days, body weight and survival rates were recorded. At 3 days post-infection (dpi), blood, spleen, and lung tissues were collected. Blood was also collected two weeks post-prime and boost via facial vein puncture. Serum was heat-inactivated, centrifuged, and stored at −80°C. Antigen-specific IgG responses were evaluated by ELISA as area under the curve.

### Ferret immunization and challenge

Ferrets for antigen groups of H3 HA monomer, 3HA-nps from mammalian 293F and insect sf9 cells were immunized in a prime-boost schedule with a three-week interval. Antigens were injected intramuscularly at a dose of 20 μg and 25 μg with AddaVax (InvivoGen) in 500 μL for prime and boost, respectively. At two weeks post-boost, all groups except for the naïve group were challenged with intranasal inoculation (6.0 Log TCID_50_/mL) of heterologous influenza H3N2 (A/Perth/16/2009) and mouse-adapted H1N1 (A/California/04/2009) viruses with a two-week interval. Nasal wash samples were collected at 1, 3, and 5 dpi of H3N2 and H1N1 challenges. The body weight and temperature were monitored for 8 dpi. Blood samples were collected at 3 weeks post-prime, 2 weeks post-boost, 2 weeks post-H3N2 challenge, and 5 days post-H1N1 challenge. A monolayer of MDCK cells (1.5 × 10^4^ cells/well) was infected with 50 μL of the 10-fold serially diluted supernatant of the nasal wash for 1 hr. Viral titers in nasal wash were measured by virus-induced cytopathic effects.

### Virus-based neutralization, ELISA, and ELISpot assays

NAb titers in sera were measured by microneutralization assay with A/Korea/01/2009 (H1N1), B/Florida/04/2006, A/Brisbane/10/2007 (H3N2), A/chicken/Korea/L433/2018 (H9N2), and A/duck/Korea/557/2016 (H7N7) influenza viruses as previously described. Antigen-specific Ab titers in sera were determined by ELISA assays with the purified antigens, H1 HA_m_-PCNA1, B HA_m_-PCNA2, and H3 HA_m_-PCNA3. ELISA data from serum IgG responses are shown as area under the curve (AUC). To measure IFN-γ secretion, a mouse IFN-γ ELISpot assay was performed using splenocytes (2×10^5^ cells/well) stimulated with 0.2 μg of PepMix^TM^ HA peptide derived from influenza A/CA/04/2009/H1N1 (JPT Peptide Technologies, Berlin, Germany).

## Author Contributions

M.S.C., M.S., and K.H.K. designed the study; H.K., S.C.M., D.B.L., J.H.J., E.J.K., J.Y.M., J.H.S., J-.L., Y.J., J.P.P.C., J.L., Y.L., Y.K.K., H.S.J., Y.H.B., X.S., J.W.L., M.S.C., M.S., and K.H.K. performed experiments, including animal experiments; J.Y.M, U.S.J., and G.L. performed TEM and AFM experiments; H.K., M.S., M.S.C., and K.H.K. wrote the paper.

## Acknowledgments

We thank Ms Jeong Suk An and Chae Yeon Lim for technical support. This work was supported by grants of the Korea Health Technology R&D Project (HV20C0054 & HV23C0060, K.H.K.) through the Korea Health Industry Development Institute (KHIDI), funded by the Ministry of Health and Welfare (MOHW), the National Research Foundation (2021R1A2C2009539, M.S.C.), and the KBRI basic research program funded by Ministry of Science and ICT (24-BR-01-03, J.Y.M.) of Korea.

