## Supplemental text for "Mosaic display of stable hemagglutinin monomers induces broad immune responses"

[1]Department of Biotechnology & Bioinformatics, Korea University, Sejong 30019, [2]College of Medicine and Medical Research Institute, Chungbuk National University, Cheongju 29644, [3]Department Food and Nutrition, Duksung Women's University, Seoul 01369, [4]Interdisciplinary Graduate Program for Artificial Intelligence Smart Convergence Technology, Korea University, Sejong 30019, Korea, [5]VIB Center for Medical Biotechnology, VIB, Ghent, Belgium, [6]Department of Biochemistry and Microbiology, Ghent University, Ghent, Belgium, [7]Research Group for Neural Circuit, Korea Brain Research Institute (KBRI), Daegu 41068, Korea, [8]Department of Engineering, Dartmouth College, Hanover, NH 03755, U.S.A.

*These authors contributed equally to this work.

**Correspondence:
 (M.S.C.)
 (M.S.)
 (K.H.K.)

Materials and methods
References
Figures S1–S12
Table S1-S2

### Materials and Methods

#### Cell lines and animals

Expi293F suspension cells derived from the 293F cell line (Life Technologies, Carlsbad, CA, USA) were cultured using Expi293 expression medium (Life Technologies) at 37℃ with 8% $CO_2$ in a shaking incubator at 150 rpm. Madin-Darby canine kidney (MDCK) cells were purchased from ATCC and grown in DMEM (Gibco BRL, Karlsruhe, Germany) supplemented with 1% penicillin-streptomycin (Gibco BRL) and 10% fetal bovine serum (Sigma-Aldrich, St. Louis, MO, USA). Five-week-old female BALB/c mice were purchased from Koatech (Pyeongtaek, Kyunggi-do, Korea), and four-month-old male ferrets were obtained from IDbio Co. Ltd. (Cheong Ju, Chungbuk, Korea).

To propagate influenza viruses, including A/Korea/01/2009 (H1N1), B/Florida/04/2006, A/Brisbane/10/2007 (H3N2), A/reassortant/X-47 (H3N2), and A/Puerto Rico/8/1934 (H1N1), MDCK cells were inoculated with the respective strains and cultured in DMEM supplemented with 1% penicillin-streptomycin at 37℃ and 5% $CO_2$. The supernatant was harvested when the cytopathic effect reached more than 80%, cleared by centrifugation at 2,000 × g for 5 min and filtered through a 0.22 µm filter to eliminate cellular debris and remaining contaminants. The filtered virus solution was purified further by centrifugation at 28,000 × g and 4℃ for 4 hr using a 30% sucrose solution in Hank's balanced salt solution. Following purification, the virus was stored at -80°C in a deep freezer, with the addition of 20% glycerol as a cryoprotectant. For influenza viruses A/duck/Korea/557/2016 (H7N7) and A/chicken/Korea/L433/2018 (H9N2), virus strains were propagated in specific pathogen-free (SPF) 10 to 11-day-old embryonated chicken eggs at 37°C for 72 hr. The eggs were chilled overnight and the allantoic fluid was harvested and stored at −80°C until needed for further experimentation.

#### Design of 3HA-np

Three hemagglutinin (HA) proteins derived from A/California/04/2009 (H1N1) (CA04), B/Florida/4/2006 (FL04), A/Gyeongnam/684/2006 (H3N2) (Gy684) influenza viruses and proliferating cell nuclear antigen (PCNA) from hyperthermophilic archaea *Saccharolobus solfataricus* were used in the design of 3HA-np. For stable HA monomer ($HA_m$) production, specific mutations were introduced into the HA residues from 11-507 for H1, 11-521 for B, and 11-498 for H3. The H1 $HA_m$ protein included the following mutations: L401S, I405S, L408S, F416E, and I419S. The B $HA_m$ protein included the mutations: L432S, I436S, L439W, V443W, L446S, T450W, L457S, and L461W. The H3 $HA_m$ protein included the mutations: V418S, V422S, L425S, V429W, and L436W. The monomeric HA mutants, H1 $HA_m$, B $HA_m$, and H3 $HA_m$, were then fused to scaffold PCNA subunits, PCNA1, PCNA2, and PCNA3_S170V, via SGG linker to produce H1 $HA_m$-PCNA1, B $HA_m$-PCNA2, and H3 $HA_m$-PCNA3, respectively. These fusion proteins were ligated to the pSecTag2A vector (Addgene, Watertown, MA, USA) using BamH1 and Not1 restriction enzymes and to the pFastBacHTA vector (Addgene) using Sal1 and Not1 restriction enzymes for mammalian and insect cell expressions, respectively. The myc and 6×His-tags were added downstream of each fusion protein. The plasmids encoding H1 $HA_m$-PCNA1, B $HA_m$-PCNA2, and H3 $HA_m$-PCNA3 were amplified in *Escherichia coli* strain DH5α and were subsequently purified using the PureLink HiPure Plasmid kit (Invitrogen, Carlsbad, CA, USA) to ensure high purity and concentration. All sequences were verified through automated sequencing conducted by Macrogen (Seoul, Korea).

### Transfection, infection, expression, and purification

To express the fusion proteins, Expi293F cells were transfected with the purified pSecTag2A vector containing the gene encoding H1 $HA_m$-PCNA1, B $HA_m$-PCNA2, or H3 $HA_m$-PCNA3 using the ExpiFectamine 293 transfection kit (Invitrogen). The transfection was performed according to the manufacturer's instructions. After transfection, the cells were incubated for 5 days at 37℃ with 8% $CO_2$ to allow for protein expression. For the insect cell expression, baculovirus containing the H1 $HA_m$-PCNA1, B $HA_m$-PCNA2, or H3 $HA_m$-PCNA3 gene was used to infect Sf9 cells. The infection was carried out at 28℃ for 3 days. After 5 days post-transfection or 3 days post-infection, the supernatants containing fusion proteins were harvested from the Expi293F or Sf9 cells, respectively, using centrifugation at 3,000 × g for 1 hr. The supernatant was collected and applied to a Ni-NTA affinity chromatography column (Qiagen, Hilden, Germany) pre-equilibrated with 20 mM Tris-HCl (pH 8.0) and 100 mM NaCl. The column was washed with an equilibration buffer containing 20 mM imidazole to remove non-specifically bound proteins. The fusion proteins were eluted from the column using an elution buffer of 20 mM Tris-HCl (pH 8.0), 100 mM NaCl, and 400 mM imidazole. The eluted protein fractions were dialyzed against 20 mM Tris-HCl (pH 8.0) and 10 mM NaCl for 16 hr at 4℃ with stirring to remove imidazole and achieve buffer exchange. Further purification was carried out using ion-exchange chromatography with a mono Q 5/50 GL column (Cytiva, Uppsala, Sweden) for each fusion protein, using a buffer system containing 20 mM Tris-HCl (pH 8.0) and 1 M NaCl. Size exclusion chromatography (SEC) was then performed using a Superdex 200 Increase 10/300 GL column (Cytiva) in Dulbecco's phosphate-buffered saline (DPBS) and Ä KTA pure system (Cytiva).

The purified H1 $HA_m$-PCNA1 and B $HA_m$-PCNA2 fusion proteins were mixed at a 1:1 molar ratio and incubated at 4℃ for 6 hr, which was then mixed with H3 $HA_m$-PCNA3 at a 1:1 ratio at 4℃ for 6 hr. The assembled 3HA-np complex was purified by SEC using a Superdex 200 Increase 10/300 GL column (Cytiva) in DPBS. To assess the purity of the protein, sodium dodecyl sulfate-polyacrylamide gel electrophoresis (SDS-PAGE) and Western blot were performed.

### SEC with multi-angle light scattering (SEC-MALS)

The shift in elution volume of the antigens upon self-assembly of the purified fusion proteins was determined by SEC on a Superdex 200 Increase 10/300 GL column (Cytiva) equilibrated with 50 mM Tris-HCl (pH 8.0) and 100 mM NaCl at a flow rate of 0.4 mL/min. We analysed the molecular weights of the antigens by means of SEC using a UFLC system (Shimadzu, Kyoto, Japan), coupled to multi-angle light scattering (SEC-MALS) using in-line WYATT-787-TS miniDAWN TREOS (Wyatt Technology, Santa Barbara, CA, USA). Purified protein (100 µL) was applied to the Superdex 200 column (Cytiva) at 0.4 mL/min, and raw data were analyzed by Astra 6 software (Wyatt Technology).

### Molecular model of 3HA-np

The overall structures of H1 HA, B HA, H3 HA, and PCNA (PDB IDs: 4EDB, 4M40, 2YP7, and 2HIK respectively) were used to construct a plausible structural model for 3HA-np, which was built manually and subjected to geometry optimization with the program Coot[1]. The amino acid sequence conservation was calculated at the ConSurf server[2]**, based on multiple sequence alignment of 150 HA proteins,** and conserved sites were presented on the structure of an H3 HA (PDB ID: 4WE8).

### Negative stained electron microscopy

Samples were prepared for electron microscopic studies by applying 3HA-np (4 μL drop at a concentration of 1.5 μM) to charged EM grids coated with carbon film. Grids were negatively stained with 1% uranyl acetate and observed under a Tecnai 20 transmission electron microscope operating at 120 kV (Thermo Fisher Scientific, Waltham, MA, USA).

### Atomic force microscopy

3HA-np in 40 μL drop at 10 ng/mL was carefully deposited onto a freshly cleaved mica surface and incubated for 15 min to allow for protein adsorption. The sample was washed ten times with distilled water, dried using $N_2$ gas, and mounted onto an atomic force microscope (AFM) scanner. The AFM data of the 3HA-np were acquired using a soft tapping mode under ambient conditions. The measurements were performed using a Multimode-VIII AFM system (Bruker, Santa Barbara, CA, USA) equipped with silicon nitride triangular AFM probes (SCANASYST-AIR, Bruker). The AFM images were acquired with a resolution of 512 × 512 pixels at a line rate of 0.3 Hz and a set point of 2 nm. The Mountains SPIP software (v9, Digital Surf, France) and NanoScope analysis software (v2.0, Bruker) were employed to analyze all the topographic information of the 3HA-np.

### Immunoblotting

The purified antigens (H1 $HA_m$-PCNA1, B $HA_m$-PCNA2, H3 $HA_m$-PCNA3, and 3HA-np) were subjected to immunoblotting by incubating with anti-6×His-tag polyclonal antibody (pAb) (ab1187, AbCam plc, Cambridge, UK), rabbit anti-influenza H1N1 (A/California/04/2009) HA monoclonal Ab (mAb) (Sino Biological, Beijing, China, 11055-RM10), anti-influenza H3N2 HA mAb (Sino Biological, 11056-RP02), or anti-influenza B HA mAb (Sino Biological, 11053-R004), diluted to a concentration of 1:1000 in TBS with 0.1% Tween 20. The antigen-Ab complexes were separated using SDS-PAGE gels and transferred to a membrane for immunoblotting analysis. For deglycosylation,

3HA-np was treated with Endo H at 10:1 ratio at 37℃ for 1 hr.

### Bio-layer interferometry

The 3HA-np's from the mammalian and insect cells were prepared at 0.5 mg/mL. Fetuin and concanavalin A (ConA) (Sigma-Aldrich, St. Louis, MO, USA, F3004 and C5275) were used in a range of 187.5–6,000 nM and 93.8–6,000 nM, respectively. The binding affinities were characterized by bio-layer interferometry (BLI) using a GatorPrime (Gator Bio, Palo Alto, CA, USA) at ambient temperature. The association step was performed by dipping the Ni-NTA biosensors with immobilized immunogen into diluted fetuin or ConA for 150 s, and dissociation was measured by inserting the biosensors back into the DPBS buffer for 120 s. Asialofetuin and mannan (Sigma-Aldrich, A4781 and M7504) were used as a control to bind to 3HA-np and an inhibitor to block the binding of 3HA-np to ConA, respectively. The mannose-rich polysaccharide, mannan, was used as a selective inhibitor of binding to ConA.

Competition assays were performed to investigate whether Abs contained in mice sera bind to different epitopes of HA. The interference signals, when the Abs in the sera were competed with the authentic mAbs, GC1517 (obtained from GC Pharma, Yongin-si, Gyeonggi-do, Korea), CR9114 (Sino Biological), and D2 H1-1/H3-1 (kindly provided by Prof. George Georgiou, University of Texas at Austin, Austin, TX, USA) against the head, stem, and interface epitopes, respectively, were detected via biolayer interferometry using GatorPrime (Gator Bio). Either trimeric or monomeric HA-coated Ni-NTA biosensor was immersed in serially diluted mAbs (0–750 μg/mL) and sera for 300 s. The biosensor was allowed to equilibrate in DPBS buffer for 120 s before proceeding to subsequent steps, and all processes involved shaking at 1,000 rpm at ambient temperature. The plots were fitted using the GatorOne software (v2.15).

### Mouse immunization and challenge

Animal procedures were performed under the approval of the Institutional Animal Care and Use Committee of Duksung Women's University (Seoul, Korea). Six-week-old BALB/c mice, after a week of adaptation, were divided into five groups: naïve, virus control, a mixture of HA monomers (H1 $HA_m$, H3 $HA_m$, and B $HA_m$), 3HA-np's from the insect and mammalian cells (the number of mice was 8–10 per group, except for the naive of 5). The mice were subjected to a prime-boost immunization with a three-week interval. The antigens were mixed with AddaVax (InvivoGen, CA, USA) at a 1:1 ratio and were intramuscularly injected into the mice at a dose of 3 μg or 5 μg of antigen in 100 μL for prime or boost, respectively. Blood samples were obtained via facial vein puncture two weeks after the prime and boost immunizations. After 3 weeks of boost immunization, mice were intranasally challenged with 5 $MLD_{50}$ of influenza virus H3N2 (A/reassortant/X-47, X47) or H1N1 (A/Puerto Rico/8/1934, PR8). Body weight and survival of the mice were recorded daily for 14 days post-infection (dpi). All naive animals remained free of any disease signs and did not experience weight loss. At 3 dpi, blood, spleen and lung tissues were collected from three mice per group.

The lung tissues were homogenized, serially diluted in DMEM, and subjected to a 10-fold serial dilution, which was then used to infect seeded monolayers of MDCK cells ($4 \times 10^5$ cells/well) in 24-well plates. Infected cells were supplemented with 1 μg/mL TPCK trypsin (Sigma-Aldrich) and 1% agarose (Lonza, Basel, Switzerland). Following incubation for 72 hr at 37℃ in a 5% $CO_2$, the plates were fixed using a 4% formaldehyde solution. Fixed plaques were stained with 0.5% crystal violet, and the number of plaques was counted.

The blood samples were allowed to coagulate at room temperature for 30 min, followed by centrifugation at 2,000 × *g* for 15 min to separate the serum. The complement factors and pathogens present in the serum were inactivated by heat treatment at 56℃ for 60 min. After centrifugation, the heat-inactivated serum samples were stored at -80℃ until further use in the study.

### Antigen-specific Ab measurement by ELISA

The evaluation of immunoglobulin G (IgG) responses was carried out using an enzyme-linked immunosorbent assay (ELISA). In this assay, 96-well ELISA plates were coated with 100 μL of 2 μg/mL H1 $HA_m$-PCNA1, B $HA_m$-PCNA2, and H3 $HA_m$-PCNA3 antigens in PBS and incubated overnight at 4℃. After blocking with 1% bovine serum albumin (BSA) in PBS containing 0.1% Tween

20 (PBS-T) for 1 hr at room temperature, the plates were washed with PBS. Serially diluted mouse sera in 100 μL, including naive mouse serum as well as a blank control (PBS-T), were added to the plate and incubated for 1 hr at room temperature. Following incubation, the plates were washed three times with PBS. Next, 100 μL of a 1:5000 dilution of goat anti-mouse IgG conjugated with horseradish peroxidase was added to each well and incubated for 1 hr at room temperature. After four additional washes with PBS, 100 μL of 3,3',5,5'-tetramethylbenzidine substrate (Sigma-Aldrich) was added to each well and incubated for 15 min at room temperature. The enzymatic reaction was stopped by adding 1 M $H_2SO_4$. To measure the absorbance at 450 nm, a Multiskan™ FC Microplate Photometer (Thermo Fisher Scientific) was used. The ELISA data from the serum IgG responses were analyzed as the area under the curve (AUC), providing a quantitative assessment of the Ab levels generated against the specific antigens.

### Virus-based neutralization assays using sera

Virus-based nAb titers were measured by microneutralization (MN) assays. MDCK cells ($1.5 \times 10^4$ cells/well) were seeded in 96-well plates for 24 hr. Influenza viruses, A/Korea/01/2009 (H1N1), B/Florida/04/2006, A/Brisbane/10/2007 (H3N2), A/chicken/Korea/L433/2018 (H9N2), and A/duck/Korea/557/2016 (H7N7) at 2 log10 $TCID_{50}$/50μL were mixed with diluted serum (in two-fold dilution from 1:4 to 1:1024) at a 1:1 ratio, which were supplemented with DMEM containing 2% FBS and 1% antibiotic-antimycotic and incubated at 37℃, 5% $CO_2$ for 1 hr. After washing the cells twice with PBS, 50 μL of a mixture containing diluted sera and influenza virus was added to the MDCK cell monolayers. After 1 hr of infection, the cells were washed twice with PBS and replaced with 150 μL MEM medium (Corning Life Science, Manassas, VA, USA) containing 1 μg/mL of TPCK trypsin. The plates were incubated for 72 hr, and the supernatant was transferred to a V-bottom 96 well microplate containing 0.5% turkey erythrocytes. Hemagglutination inhibition assays were performed at different concentrations of sera.

### Ferret immunization and challenge

Ferret experiments were approved by the Institutional Animal Care and Use Committee of Chungbuk National University (Chungbuk, Korea). Four-month-old ferrets were confirmed to be seronegative for seasonal H1N1 and H3N2-like viruses through hemagglutination inhibition assays prior to the experiments. They were divided into five groups of the naïve, virus control, H3 HA monomer, 3HA-nps from mammalian and insect cells (the number of ferrets was 4–5 per group, except for the naive of 2). Ferrets of antigen groups were given with two intramuscular immunizations with a 3-week interval at a dose of 20 μg or 25 μg of antigen with AddaVax in 500 μL for prime or boost, respectively. Ferrets of the naïve and virus control groups were injected intramuscularly with 500 μL of PBS in AddaVax and only PBS, respectively, at a 3-week interval. Blood samples were collected at 3 weeks after prime and at 2 weeks after boost immunization.

The immunized ferrets were lightly anesthetized with alfaxalone and xylazine two weeks after receiving the booster and intranasally inoculated with 6.0 Log10 $TCID_{50}$/mL of H3N2 virus (A/Perth/16/2009). At 14 dpi, blood samples were collected from all H3N2-infected ferrets and the ferrets were inoculated intranasally with 6.0 Log10 $TCID_{50}$ of mouse-adapted H1N1 (A/California/04/2009) virus. Nasal wash samples were collected at 1, 3, and 5 dpi of H3N2 and H1N1 infections. The body weight and temperature were recorded for 8 dpi. At 5 dpi of H1N1 infection, blood and spleen tissues were collected from two ferrets in each group, and the tissues were stored at -80℃ until further use. A monolayer of MDCK cells ($1.5 \times 10^4$ cells/well) was incubated in 96-well plates for 24 hr before infection. Cell monolayers were washed two times with PBS and infected with 50 μL of the 10-fold serially diluted supernatant of the nasal wash for 1 hr. After infection, cell monolayers were washed twice with PBS and replaced 150 μL MEM medium with 1 μg/mL TPCK-trypsin. After 72 hr of infection, the supernatant determined the viral replication by $TCID_{50}$ assay.

#### IFN-γ ELISpot assay

Antigen-specific IFN-γ secretions were assessed by ELISpot assay using an IFN-γ ELISpot kit (Mabtech, Nacka Strand, Sweden). Briefly, splenocytes from challenged mouse or ferret ($2.5 \times 10^5$ cells/well) were stimulated with 0.2 μg of PepMix™ HA peptide, a peptide pool of 15-mer sequences with 11 amino acid overlap covering the complete sequence of HA derived from influenza A/CA/04/2009/H1N1 (JPT Peptide Technologies, Berlin, Germany) for 24 h at 37°C in a 5% $CO_2$. Splenotyes without stimulation with the peptides were used as a control. Cells were washed with DPBS (Welgene, Gyeongan, Korea) and incubated with the detection anti-mouse IFN-γ mAb (R4-6A2, Mabtech, Cincinnati, USA) to 1 μg/mL in PBS containing 0.5% fetal calf serum (FCS) for 2 hrs at room temperature. The plates were washed 5 times with PBS and treated with streptavidin-ALP diluted 1:1000 in PBS containing 0.5% FCS for 1 hr at room temperature. After washing, each well was incubated with BCIP/NBT-plus substrate solution for 10-30 min at room temperature. The reaction was stopped by washing with water. The resulting spots were manually counted, and the mean spot counts of negative control wells were to generate normalized readings. Background response levels were determined using splenocytes taken from the naive animal.

### Supplementary figure legends

**Fig. S1. Design of the HA-PCNA constructs for insect cell expression and mutation sites on HA structures.** (Upper panel) Design of the 3HA-np components, H1 $HA_m$-PCNA1, B $HA_m$-PCNA2, and H3 $HA_m$-PCNA3, showing schematic diagrams of influenza virus HA-PCNA fusion proteins. The mutation sites for stable HA monomer ($HA_m$) are shown in red characters and HA structures (lower panel). The HA regions are derived from 11 to 507 for H1, 11 to 521 for B, and 11 to 498 for H3 HA proteins, which are fused to the N-terminus of PCNA via SGG linker. The 6×His-tag is fused to the N-terminus of the gene for insect cell expression.

**Fig. S2. Mutation sites for HA mutant monomers.** The mutation sites at the monomer-monomer interface (left panel) and for disulfide bond formation (right panel) for HA monomers, H1 HA, B HA, and H3 HA are highlighted in spheres in the structure of HA monomer. An HA trimer structure is shown with interface mutation sites are highlighted in spheres with side and top views (in the box).

**Fig. S3. Purification and assembly of the mosaic 3HA-np components.** (A) Purification of H1 $HA_m$-PCNA1, B $HA_m$-PCNA2, and H3 $HA_m$-PCNA3 in the mammalian (left panel) and insect (right panel) cells. The elution profiles from the final SEC step, using Superdex 200 increase 10/300 GL, and SDS-PAGE results are shown. (B) The shifts in elution profiles upon self-assembly of the fusion protein components are shown for comparison, using SEC on Superdex 200 increase 10/300 GL. The purified H1 $HA_m$-PCNA1and B $HA_m$-PCNA2 were mixed in a 1:1 molar ratio, to which H3 $HA_m$-PCNA3 was added in a molar ratio of 1:1.

**Fig. S4. Characterization of 3HA-np using SEC.** The elution profile of 3HA-nps of mammalian (left panel) and insect (right panel) cells from the final SEC step, using Superdex 200 increase 10/300 GL, and SDS-PAGE results are also shown.

**Fig. S5. Characterization of the components of 3HA-np using SEC-MALS.** The purified H1 $HA_m$-PCNA1, B $HA_m$-PCNA2, and H3 $HA_m$-PCNA3 are characterized using SEC-MALS to show molecular weights of 91.3 kDa (±12.2%), 91.0 kDa (±1.8%), and 103.5 kDa (±1.6%) in the mammalian cell (left panel) and 82.0 kDa (±12.2%), 92.4 kDa (±1.8%), and 96.1 kDa (±1.6%) in the insect cell (right panel), respectively.

**Fig. S6. Binding affinity of 3HA-np to fetuin.** Determination of binding affinities between 3HA-np and fetuin was performed using biolayer interferometry. Analysis of fetuin (in the range of 187.5 ~ 6,000 nM) binding to immobilized 3HA-np was done with Ni-NTA biosensors. The dissociation constant ($K_D$) was $1.42 \times 10^{-6}$ M and $1.47 \times 10^{-6}$ M in the 3HA-np from mammalian (upper panel) and insect (lower panel) cell expression, respectively.

**Fig. S7. Effects of glycosylation on binding of 3HA-np to ConA.** Binding of 3HA-np from the mammalian and insect cell expression to ConA were evaluated upon deglycosylation by Endo H using biolayer interferometry. ConA, a mannose binder, binding was performed using Ni-NTA biosensors immobilized with 3HA-np. Binding of 3HA-np's to ConA, a mannose binder, was inhibited by treatment with Endo H, which was further reduced by the presence of mannan, a mannose-rich polysaccharide and inhibitor of binding to ConA.

**Fig. S8. Stability of the 3HA-np components**. Time-course experiments of H1 $HA_m$-PCNA1, B $HA_m$-PCNA2, and H3 $HA_m$-PCNA3 monitored at –80℃, 4℃, 25℃, and 37℃ over a 28-day storage period (0, 1, 4, 7, 14, and 28 days), using SDS-PAGE for the protein components expressed in the mammalian (upper panel) and insect (lower panel) cells, respectively.

**Fig. S9. Competition assays for Abs in mice sera binding to epitopes of HA.** The H1 HA trimer (left panel) or HA monomer (right panel) was loaded onto the Ni-NTA biosensor, and mAbs GC1517, CR9114, and D2 H1-1/H3-1 directed against spatially distinct epitopes, the head, stem, and interface, were used, respectively. The interference signals, when the Abs in the sera were competed with the authentic mAb, were detected via biolayer interferometry.

**Fig. S10. Binding of mAbs to H1 and H3 $HA_m$s.** Binding of D2 H1-1/H3-1, CR9114, and GC1517 to H1 and H3 HA monomers were evaluated using biolayer interferometry. The H1 or H3 $HA_m$ monomers were loaded onto the Ni-NTA biosensor, and mAbs GC1517, CR9114, and D2 H1-1/H3-1 were evaluated in the range of 0–20 μM.

**Fig. S11. Characterization of cell-mediated immune response using IFN-γ ELISpot assay**. IFN-γ responses (upper panel) and averaged ELISpot results in spot forming units (SFUs) per $2 \times 10^5$ splenocytes (lower panel) isolated from mice at 3 dpi of H3N2 or H1N1 challenge. The splenocytes were stimulated with 0.2 mg of PepMix™ HA peptide derived from influenza A/CA/04/2009/H1N1 to evaluate antigen-specific IFN-γ-producing T-cells using an ELISpot assay. The samples were analyzed in duplicates.

**Fig. S12. Representative IFN-γ ELISpot results of ferret sera**. IFN-γ responses (upper panel) and averaged ELISpot results in spot forming units (SFUs) per $2 \times 10^5$ splenocytes (lower panel) isolated from ferrets. Ferrets were challenged with heterologous influenza H3N2 and maH1N1 viruses at a 2-week interval. The spleen was collected from two ferrets in each group at 5 dpi of the H1N1 challenge. The splenocytes were stimulated with 0.2 mg of PepMix™ HA peptide derived from influenza A/CA/04/2009/H1N1 to evaluate antigen-specific IFN-γ-producing T-cells using an ELISpot assay. The samples were analyzed in duplicates.

**Table S1. List of primers used in this study**

| HA-PCNA | | Primer sequences (5' - 3') |
|---|---|---|
| H1 | for | ACGCGTCGACTCCGGGGACACATTATGTATAGG |
| | rev | GAACATACCTCCGGATTCTCTGTTTAATTTTGCTTCC |
| PCNA1 | for | GAGAATCCGGAGGTATGTTCAAGATCGTGTACCCCAAC |
| | rev | ATAGTTTAGCGGCCGCTTAACCACGGGGAGCGATCCAG |
| B | for | ACGCGTCGACTCCTCCGATCGTATCTGCACTGGCATC |
| | rev | CATCATACCTCCGGAACCAGTGGGGAGAGAAAATTCTCC |
| PCNA2 | for | CCACTGGTTCCGGAGGTATGATGAAGGCTAAGGTGATC |
| | rev | ATAGTTTAGCGGCCGCTTAGTCAGCACGGGGAGCGATG |
| H3 | for | ACGCGTCGACGCGGATCCCGGGGCAACGCTG |
| | rev | CTTCATACCTCCGGAAGCCCGGTTGTTTAATGCTTCATC |
| PCNA3 | for | CCGGGCTTCCGGAGGTATGAAGGTGGTGTACGACGACG |
| | rev | ATAGTTTAGCGGCCGCTTAACCCTTGGGAGCCAGCAGGTAG |

**Table S2. List of amino acid sequences for HA$_m$-PCNA**

| Name | |
|---|---|
| HI HA$_m$-PCNA1 | METDTLLLWVLLLWVPGSTGDAAQPARRAVRSLVPSSDPLQCGGILQSGDTLCIGYHANNSTD |
| (mammalian) | TVDTCLEKNVTVTHSVNLLEDKHNGKLCKLRGVAPLHLGKCNIAGWILGNPECESLSTASSWS |
| | YIVETPSSDNGTCYPGDFIDYEELREQLSSVSSFERFEIFPKTSSWPNHDSNKGVTAACPHAGAKS |
| | FYKNLIWLVKKGNSYPKLSKSYINDKGKEVLVLWGIHHPSTSADQQSLYQNADAYVFVGSSRY |
| | SKKFKPEIAIRPKVRDQEGRMNYYWTLVEPGDKITFEATGNLVVPRYAFAMERNAGSGIIISDTP |
| | VHDCNTTCQTPKGAINTSLPFQNIHPITIGKCPKYVKSTKLRLATGCRNIPSIQSRGLFGAIAGFIE |
| | GGWTGMVDGWYGYHHQNEQGSGYAADLKSTQNAIDEITNKVNSVIEKMNTQFTAVGKEFNH |
| | SEKRSENSNKKVDDGELDWWTYNAELLVLLENCRTLDYCDSNVKNLYEKVRSQLKNNAKEIG |
| | NGCFEFYHKCDNTCMESVKNGTYDYPKYSEEAKLNRESGGMFKIVYPNAKDFFSFINSITNVTD |
| | SIILNFTEDGIFSRHLTEDKVLMAIMRIPKDVLSEYSIDSPTSVKLDVSSVKKILSKASSKKATIELT |
| | ETDSGLKIIIRDEKSGAKSKIKIKAEKGQVEQLTEPKVNLAVNFTTDESVLNVIAADVTLVGEEM |
| | RISTEEDKIKIEAGEEGKRYVAFLMKDKPLKELSIDTSASSSYSAEMFKDAVKGLRGFSAPTMVS |
| | FGENLPMKIDVEAVSGGHMIFWIAPRGAAARGGPEQKLISEEDLNSAVDHHHHHH |
| HI HA$_m$-PCNA1 (insect) | MSYYHHHHHHDYDIPTTENLYFQGVDSGDTLCIGYHANNSTDTVDTCLEKNVTVTHSVNLLED |
| | KHNGKLCKLRGVAPLHLGKCNIAGWILGNPECESLSTASSWSYIVETPSSDNGTCYPGDFIDYEE |
| | LREQLSSVSSFERFEIFPKTSSWPNHDSNKGVTAACPHAGAKSFYKNLIWLVKKGNSYPKLSKS |
| | YINDKGKEVLVLWGIHHPSTSADQQSLYQNADAYVFVGSSRYSKKFKPEIAIRPKVRDQEGRM |
| | NYYWTLVEPGDKITFEATGNLVVPRYAFAMERNAGSGIIISDTPVHDCNTTCQTPKGAINTSLPF |
| | QNIHPITIGKCPKYVKSTKLRLATGCRNIPSIQSRGLFGAIAGFIEGGWTGMVDGWYGYHHQNE |
| | QGSGYAADLKSTQNAIDEITNKVNSVIEKMNTQFTAVGKEFNHSEKRSENSNKKVDDGELDW |
| | WTYNAELLVLLENCRTLDYCDSNVKNLYEKVRSQLKNNAKEIGNGCFEFYHKCDNTCMESVK |
| | NGTYDYPKYSEEAKLNRESGGMFKIVYPNAKDFFSFINSITNVTDSIILNFTEDGIFSRHLTEDKV |
| | LMAIMRIPKDVLSEYSIDSPTSVKLDVSSVKKILSKASSKKATIELTETDSGLKIIIRDEKSGAKSKI |
| | KIKAEKGQVEQLTEPKVNLAVNFTTDESVLNVIAADVTLVGEEMRISTEEDKIKIEAGEEGKRY |
| | VAFLMKDKPLKELSIDTSASSSYSAEMFKDAVKGLRGFSAPTMVSFGENLPMKIDVEAVSGGH |
| | MIFWIAPRG |

| | |
|---|---|
| B HA$_m$-PCNA2 (mammalian) | METDTLLLWVLLLWVPGSTGDAAQPARRAVRSLVPSSDPLQCGGILQSSDRICTGITSSNSPHV VKTATQGEVNVTGVIPLTTTPTKSYFANLKGTRTRGKLCPDCLNCTDLDVALGRPMCVGTTPS AKASILHEVRPVTSGCFPIMHDRTKIRQLPNLLRGYENIRLSTQNVIDAEKAPGGPYRLGTSGSCP NATSKIGFFATMAWAVPKDNYKNATNPLTVEVPYICTEGEDQITVWGFHSDNKTQMKNLYGD SNPQKFTSSANGVTTHYVSQIGDFPDQTEDGGLPQSGRIVVDYMMQKPGKTGTIVYQRGVLLP QKVWCASGRSKVIKGSLPLIGEADCLHEKYGGLNKSKPYYTGEHAKAIGNCPIWVKTPLKLAN GTKYRPPAKLLKERGFFGAIAGFLEGGWEGMIAGWHGYTSHGAHGVAVAADLKSTQEAINKIT KNLNSLSELEVKNLQRLSGAMDESHNESLEWDEKWDDSRADWISSQIESAVLWSNEGIINSEDE HLLALERKLKKMLGPSAVDIGNGCFETKHKCNQTCLDRIAAGTFNAGEFSLPSGGMMKAKVID AVSFSYILRTVGDFLSEANFIVTKEGIRVSGIDPSRVVFLDIFLPSSYFEGFEVSQEKEIIGFKLEDV NDILKRVLKDDTLILSSNESKLTLTFDGEFTRSFELPLIQVESTQPPSVNLEFPFKAQLLTITFADII DELSDLGEVLNIHSKENKLYFEVIGDLVTVKVELSTDNGTLLEASGADVSSSYGMEYVANTTK MRRASDSMELYFGSQIPLKLRFKLPQEGYGDFYIAPRADAAARGGPEQKLISEEDLNSAVDHHH HHH |
| B HA$_m$-PCNA2 (insect) | MSYYHHHHHHDYDIPTTENLYFQGVDSSDRICTGITSSNSPHVVKTATQGEVNVTGVIPLTTTPT KSYFANLKGTRTRGKLCPDCLNCTDLDVALGRPMCVGTTPSAKASILHEVRPVTSGCFPIMHDR TKIRQLPNLLRGYENIRLSTQNVIDAEKAPGGPYRLGTSGSCPNATSKIGFFATMAWAVPKDNY KNATNPLTVEVPYICTEGEDQITVWGFHSDNKTQMKNLYGDSNPQKFTSSANGVTTHYVSQIG DFPDQTEDGGLPQSGRIVVDYMMQKPGKTGTIVYQRGVLLPQKVWCASGRSKVIKGSLPLIGE ADCLHEKYGGLNKSKPYYTGEHAKAIGNCPIWVKTPLKLANGTKYRPPAKLLKERGFFGAIAG FLEGGWEGMIAGWHGYTSHGAHGVAVAADLKSTQEAINKITKNLNSLSELEVKNLQRLSGAM DESHNESLEWDEKWDDSRADWISSQIESAVLWSNEGIINSEDEHLLALERKLKKMLGPSAVDIG NGCFETKHKCNQTCLDRIAAGTFNAGEFSLPSGGMMKAKVIDAVSFSYILRTVGDFLSEANFIV TKEGIRVSGIDPSRVVFLDIFLPSSYFEGFEVSQEKEIIGFKLEDVNDILKRVLKDDTLILSSNESKL TLTFDGEFTRSFELPLIQVESTQPPSVNLEFPFKAQLLTITFADIIDELSDLGEVLNIHSKENKLYFE VIGDLVTVKVELSTDNGTLLEASGADVSSSYGMEYVANTTKMRRASDSMELYFGSQIPLKLRF KLPQEGYGDFYIAPRAD |

| | |
|---|---|
| H3 $HA_m$-PCNA3 | METDTLLLWVLLLWVPGSTGDAAQPARRAVRSLGTADPGATLCLGHHAVQNGTIVKTITNDQI |
| (mammalian) | EVTNATELVQNSSTGGICDSPHQILDGENCTLIDALLGDPQCDGFQNKKWDLFVERSKAYSNCY |
| | PYDVPDYASLRSLVASSGTLEFNNESFNWTGVTQNGTSSACKRGSNNSFFSRLNWLTHSKFKYP |
| | ALNVTMPNNEEFDKLYIWGVHHPGTDNDQIFLYAQASGRITVSTKRSQQTVIPNIGSRPRVRDIP |
| | SRISIYWTIVKPGDILLINSTGNLIAPRGYFKIRSGKSSIMRSDAPIGKCNSECITPNGSIPNDKPFQN |
| | VNRITYGACPRYVKQNTLKLATGCRNVPEKQTRGIFGAIAGFIENGWEGMVDGWYGFRHQNS |
| | EGIGQAADLKSTQAAIDQINGKLNRLIGKTNEKFHQIEKEFSESEGRSQDSEKYWEDTKIDWWS |
| | YNAELLVALENQHTIDLCDSEMNKLFEKTKKQLRENAEDMGNGCFKIYHKCDNACIGSIRNGT |
| | YDHDVYRDEALNNRSGGMKVVYDDVRVLKDIIQALARLVDEAVLKFKQDSVELVALDRAHIS |
| | LISVNLPREMFKEYDVNDEFKFGFNTQFLMKILKVAKRKEAIEIASESPDSVIINIIGSTNREFNVR |
| | NLEVSEQEIPEINLQFDISATISSDGFKSAISEVSTVTDNVVVEGHEDRILIKAEGEVEVEVEFSKD |
| | TGGLQDLEFSKESKNSYSAEYLDDVLSLTKLSDYVKISFGNQKPLQLFFNMEGGGKVTYLLAPK |
| | GAAARGGPEQKLISEEDLNSAVDHHHHHH |
| H3 $HA_m$-PCNA3 (insect) | MSYYHHHHHHDYDIPTTENLYFQGVDADPGATLCLGHHAVQNGTIVKTITNDQIEVTNATELV |
| | QNSSTGGICDSPHQILDGENCTLIDALLGDPQCDGFQNKKWDLFVERSKAYSNCYPYDVPDYAS |
| | LRSLVASSGTLEFNNESFNWTGVTQNGTSSACKRGSNNSFFSRLNWLTHSKFKYPALNVTMPN |
| | NEEFDKLYIWGVHHPGTDNDQIFLYAQASGRITVSTKRSQQTVIPNIGSRPRVRDIPSRISIYWTIV |
| | KPGDILLINSTGNLIAPRGYFKIRSGKSSIMRSDAPIGKCNSECITPNGSIPNDKPFQNVNRITYGA |
| | CPRYVKQNTLKLATGCRNVPEKQTRGIFGAIAGFIENGWEGMVDGWYGFRHQNSEGIGQAAD |
| | LKSTQAAIDQINGKLNRLIGKTNEKFHQIEKEFSESEGRSQDSEKYWEDTKIDWWSYNAELLVA |
| | LENQHTIDLCDSEMNKLFEKTKKQLRENAEDMGNGCFKIYHKCDNACIGSIRNGTYDHDVYRD |
| | EALNNRSGGMKVVYDDVRVLKDIIQALARLVDEAVLKFKQDSVELVALDRAHISLISVNLPRE |
| | MFKEYDVNDEFKFGFNTQFLMKILKVAKRKEAIEIASESPDSVIINIIGSTNREFNVRNLEVSEQEI |
| | PEINLQFDISATISSDGFKSAISEVSTVTDNVVVEGHEDRILIKAEGEVEVEVEFSKDTGGLQDLEF |
| | SKESKNSYSAEYLDDVLSLTKLSDYVKISFGNQKPLQLFFNMEGGGKVTYLLAPKG |
