## Supplemental Figure for "Mosaic display of stable hemagglutinin monomers induces broad immune responses"

### Construct design for Insect cell expression

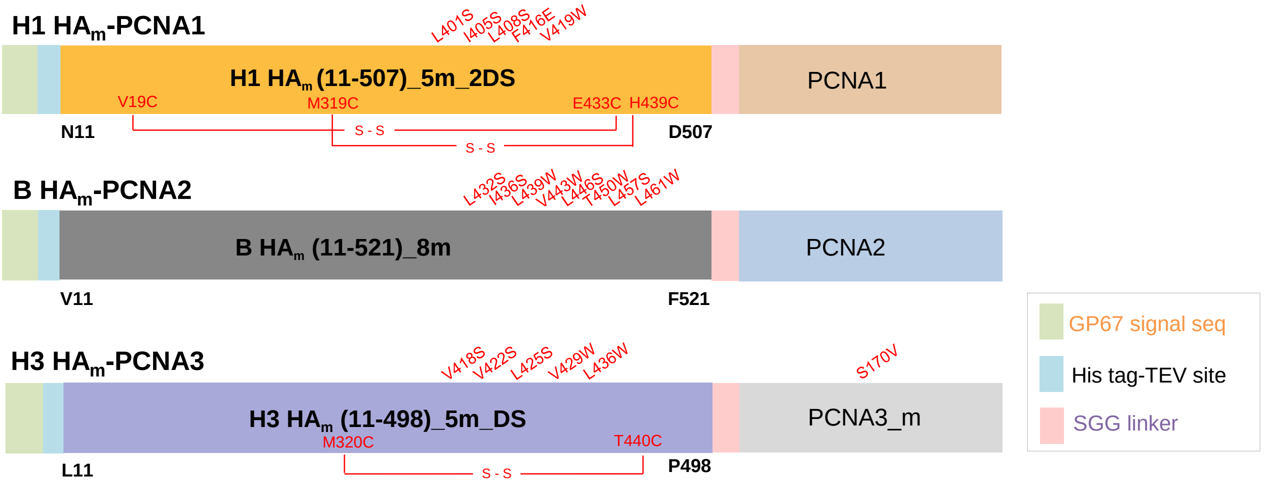


### Supplementary Fig. S1

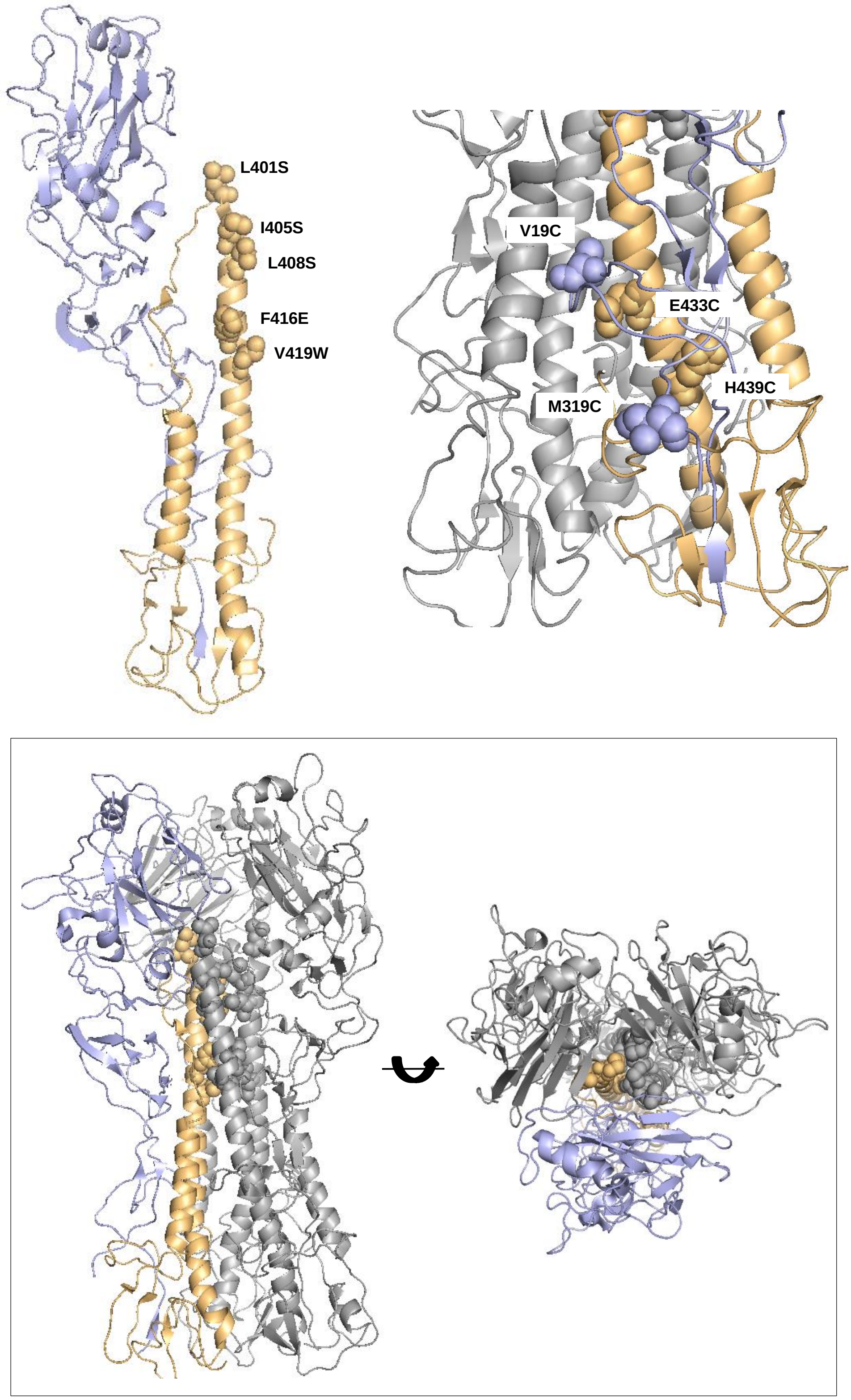


### Supplementary Fig. S2

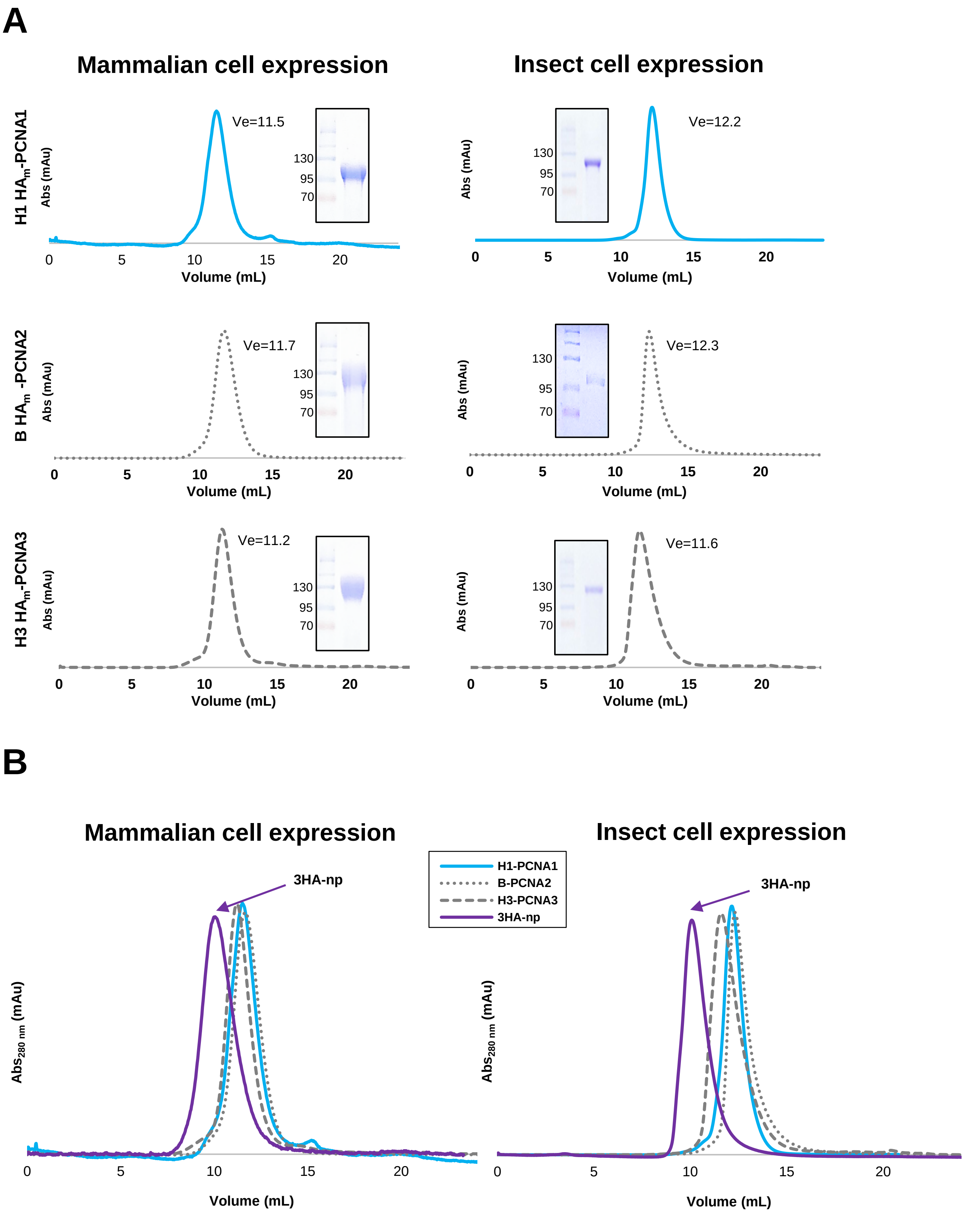


### Supplementary Fig. S3

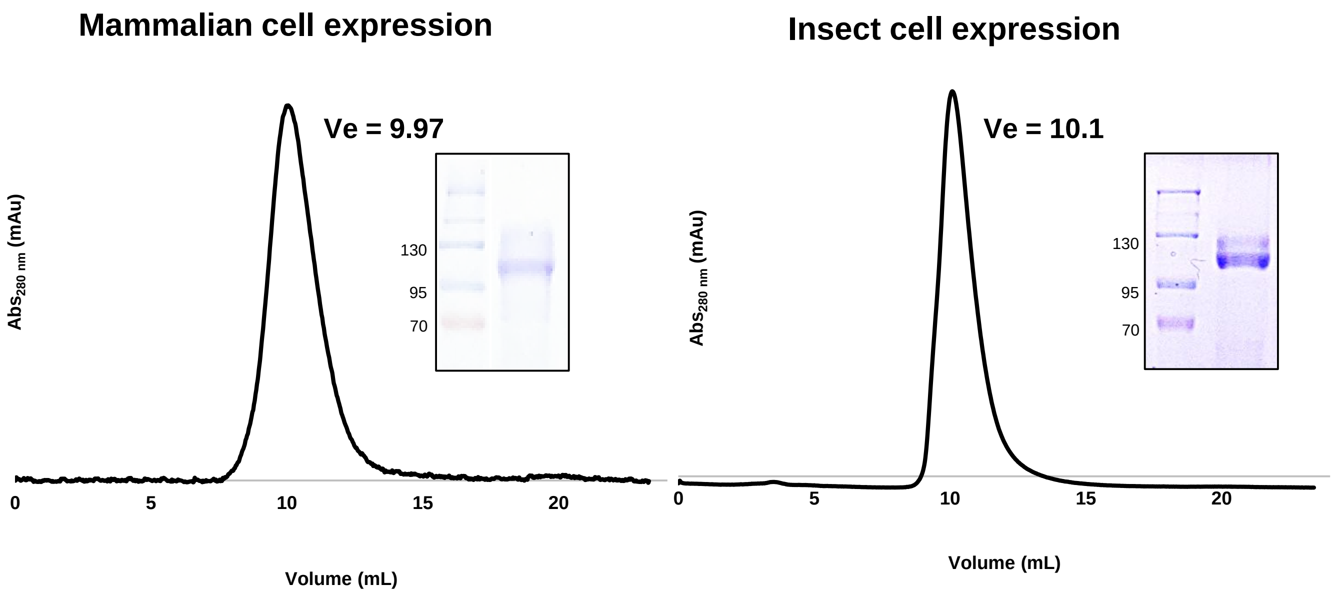


### Supplementary Fig. S4

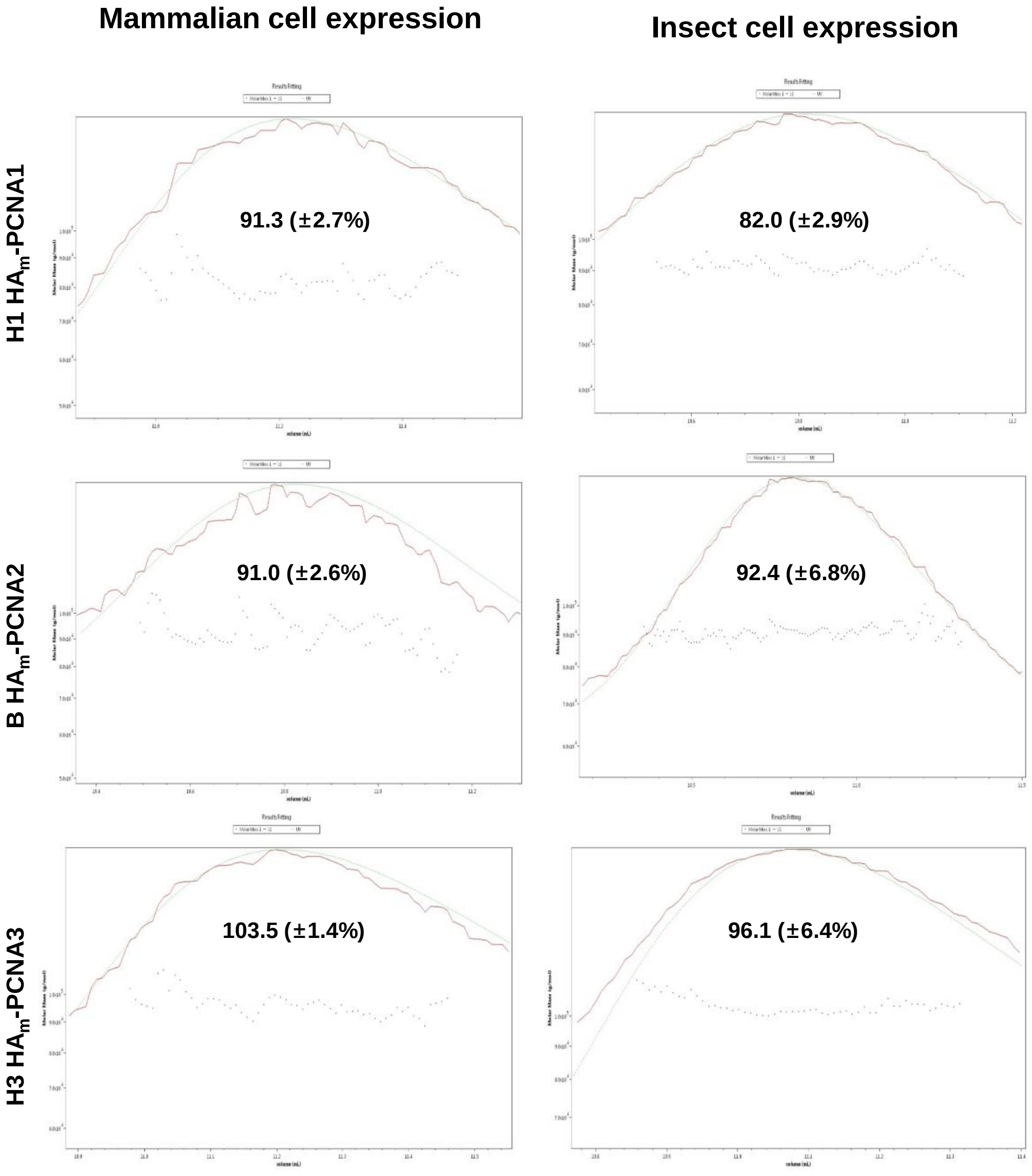


### Supplementary Fig. S5

**3HA-np (mammalian)**

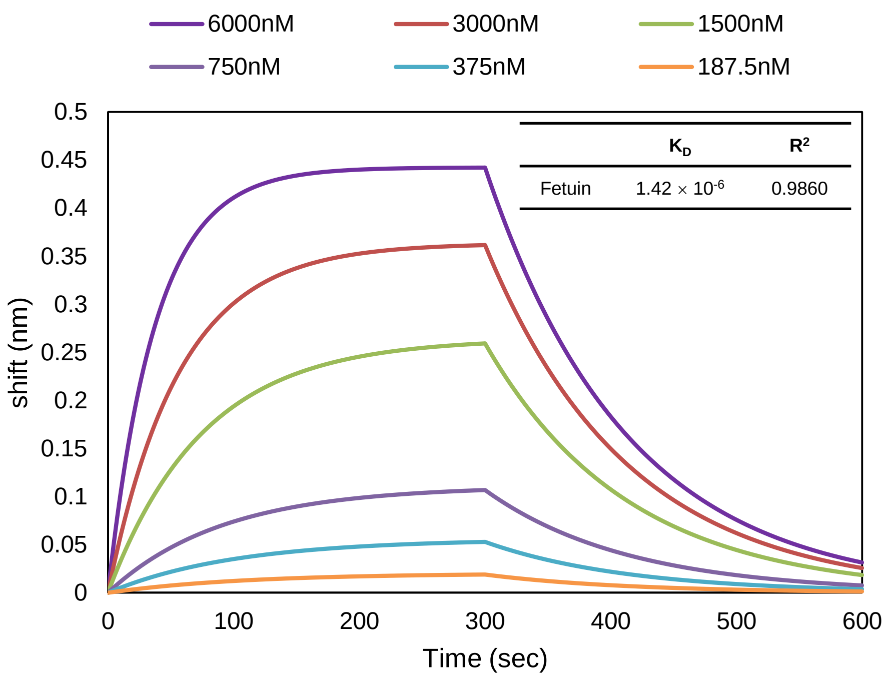


**3HA-np (insect)**

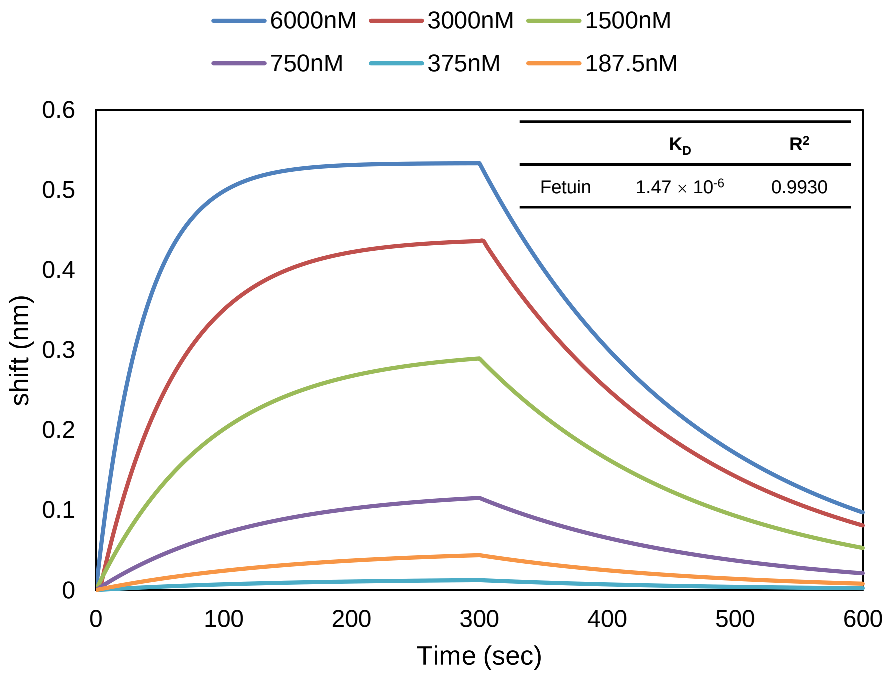


### Supplementary Fig. S6

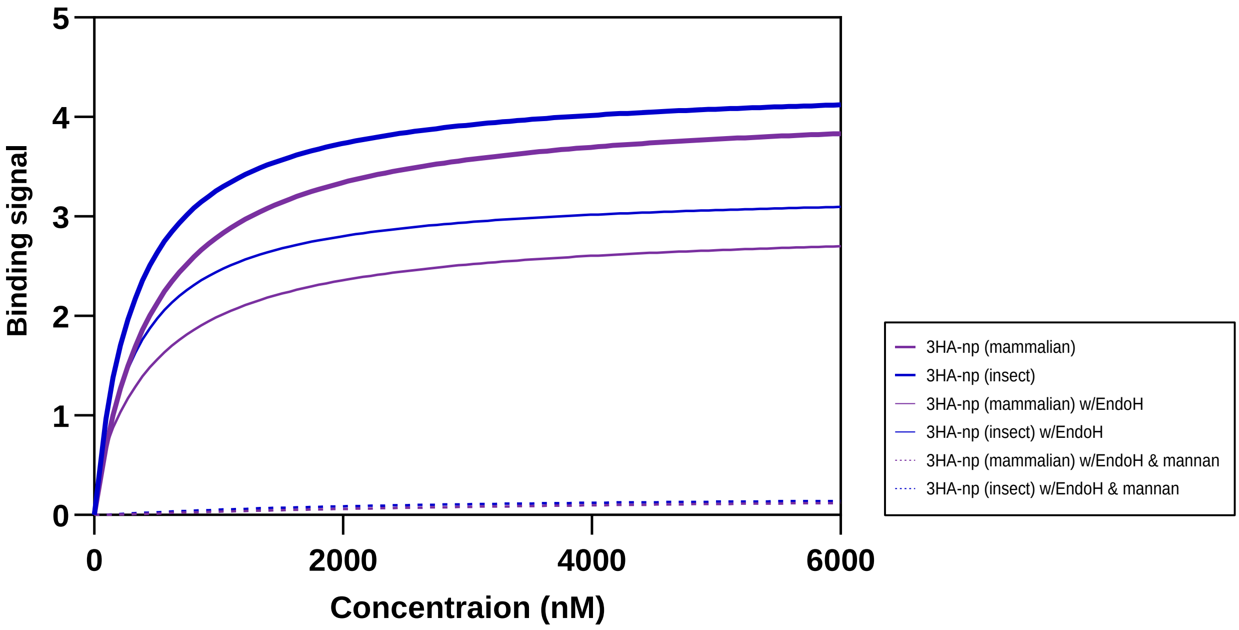


### Supplementary Fig. S7

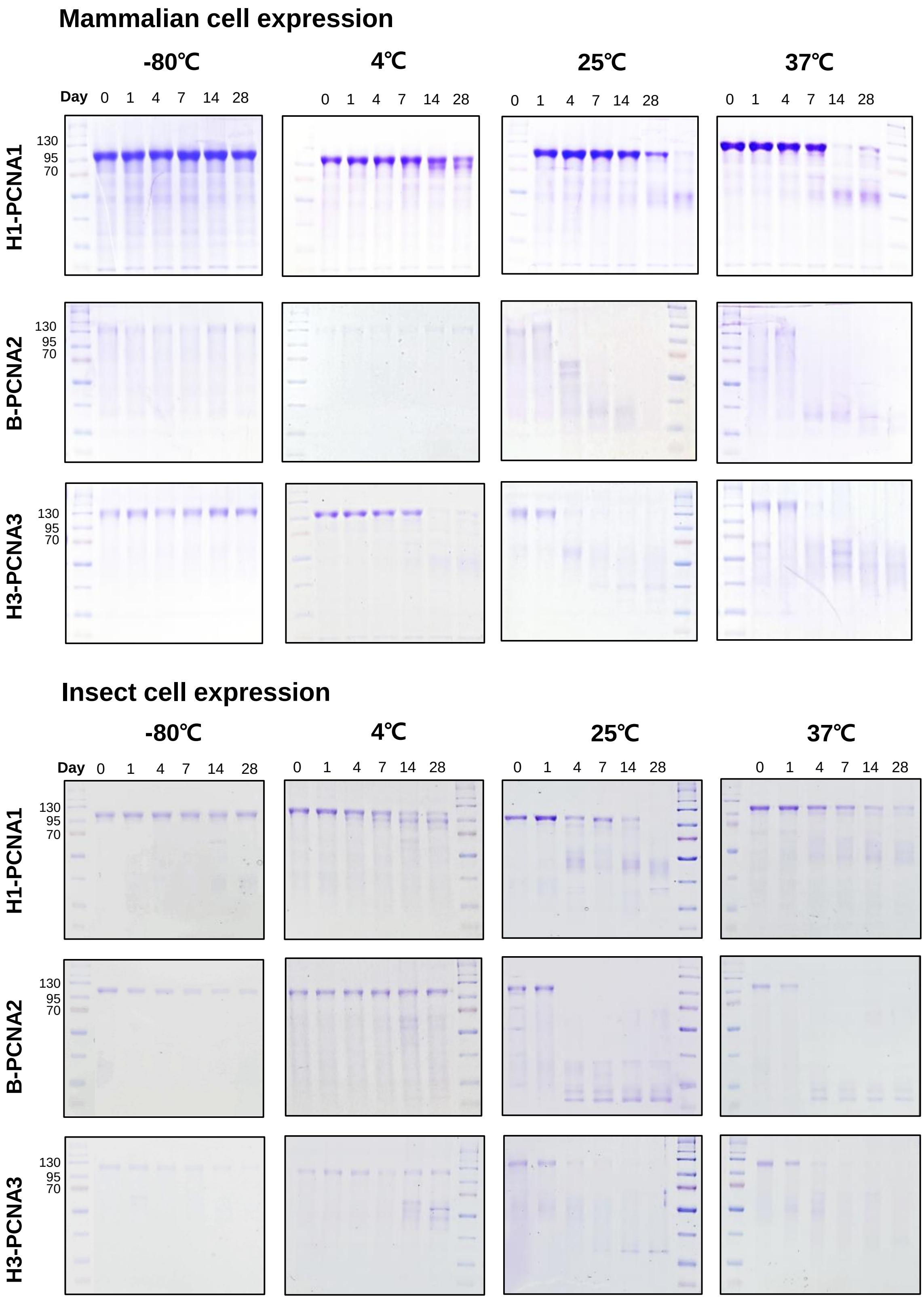


### Supplementary Fig. S8

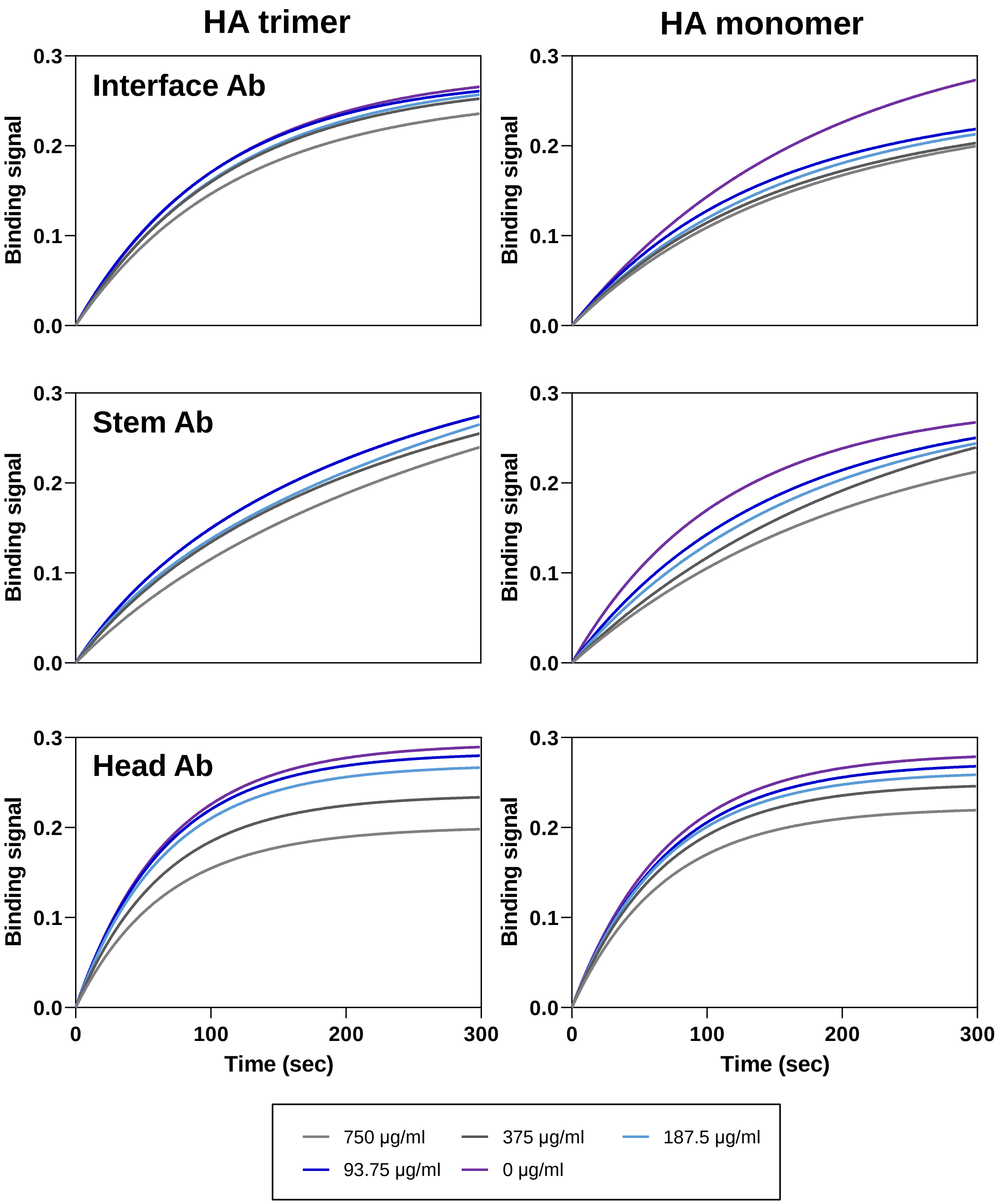


### Supplementary Fig. S9

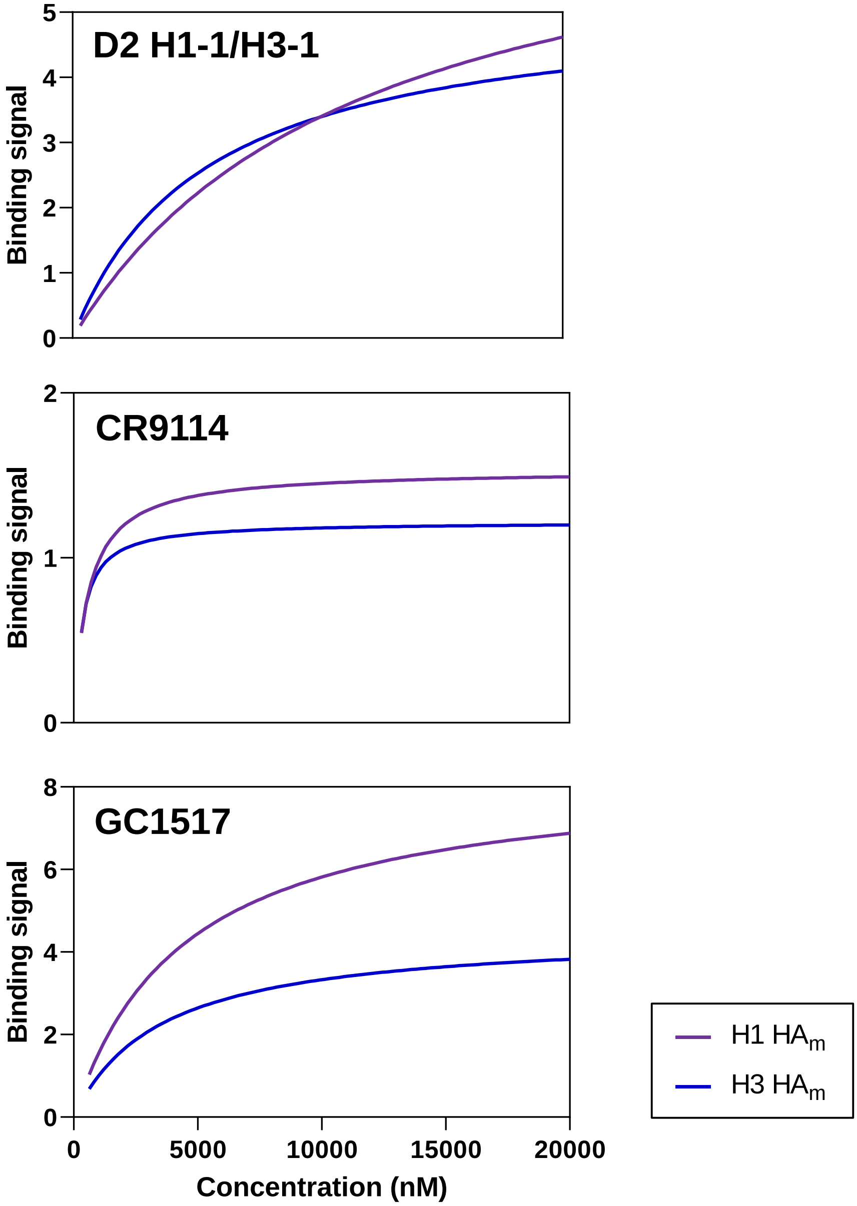


### Supplementary Fig. S10

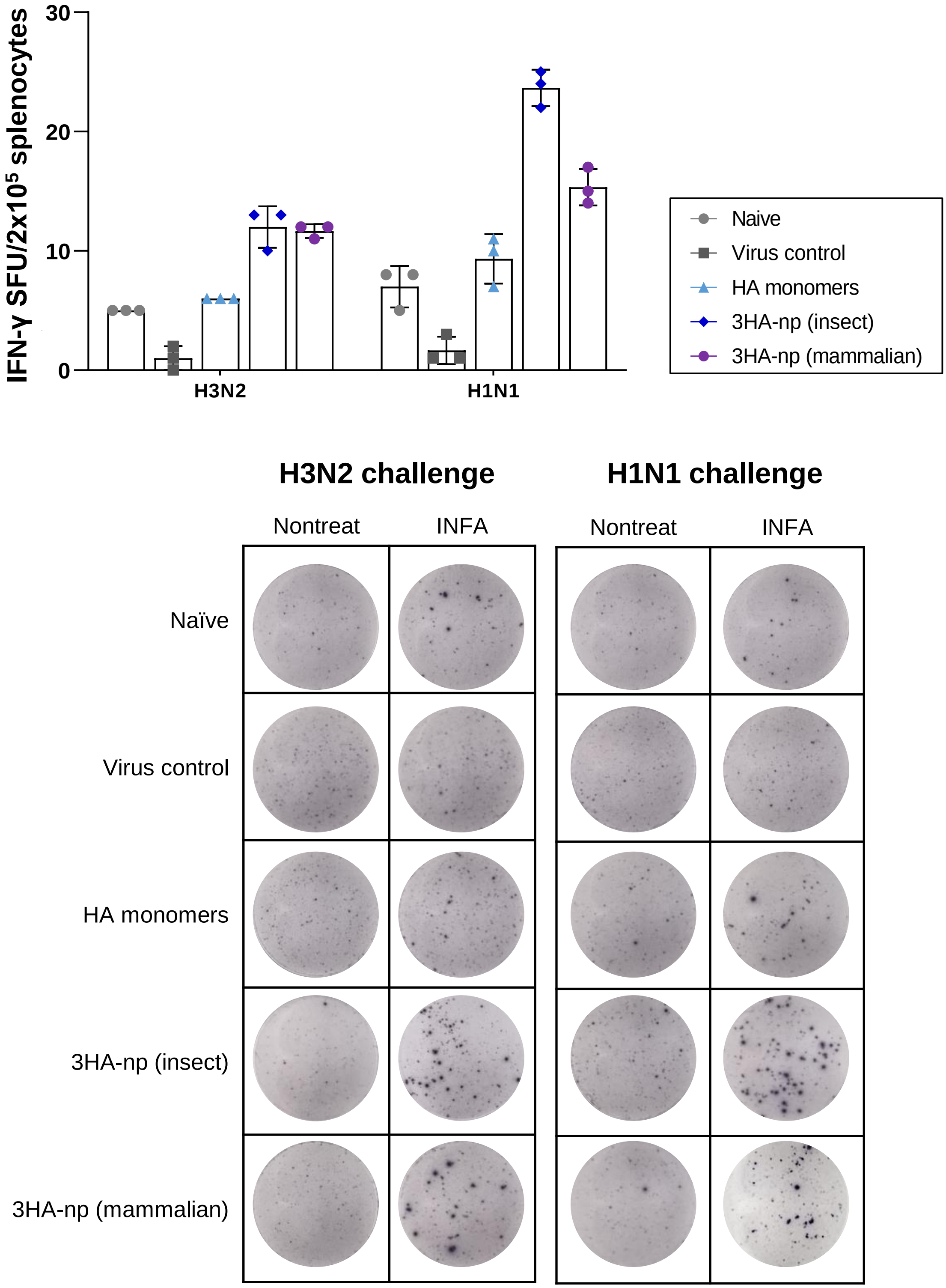


### Supplementary Fig. S11

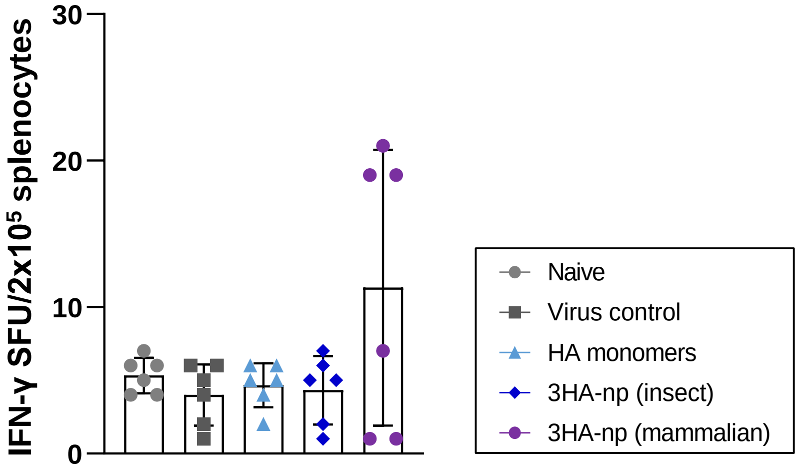


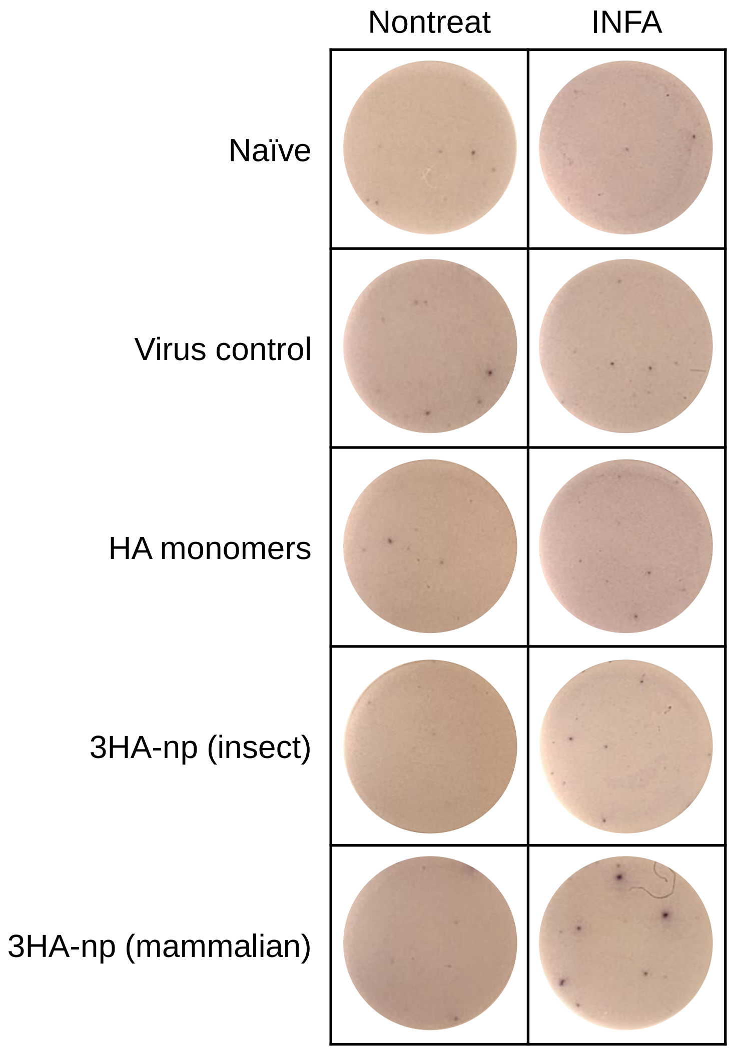


### Supplementary Fig. S12
